# A Challenge To The Assumption That Short- versus Long-Access Groups of Opioid Users Represent Distinct Phenotypes

**DOI:** 10.1101/2025.11.10.687622

**Authors:** Michael A. Giunta, Indu Mithra Madhuranthakam, Martin O. Job

## Abstract

**Rationale:** One of the current models for drug (opioid) user typology employs differential access conditions to categorize takers into short-access and long-access groups (ShA, LgA) with the rationale that these groups represent distinct opioid user phenotypes. However, with the idea that differential vulnerabilities to opioid effects are already present prior to experimenter assignment into these groups, it is unclear that these groups represent distinct opioid user types. To clarify this, we have developed a method that includes principal component analysis-gaussian mixtures model clustering of variables derived from a new CENTERED (<u>C</u>umulative <u>E</u>xperience-<u>N</u>ormalized <u>T</u>ime-<u>E</u>ffect on <u>R</u>esponse as an <u>E</u>xponential <u>D</u>ecay structure) model. The goal of this study was to utilize CENTERED clustering to test the hypothesis that ShA and LgA groups defined by the experimenter via random assignment are composed of mixtures of individuals that belong to distinct opioid user types.

**Methods:** We reanalyzed data from a previous study in which the experimenter assigned male Sprague Dawley rats (n = 30) self-administering 0.1 mg/kg/infusion oxycodone for 20 days into ShA (3h-access) and LgA (9h-access). We conducted CENTERED clustering on all takers, irrespective of assigned group(s) to determine if the experimenter-assigned groups included mixtures of individuals from groups (opioid user types) identified via CENTERED clustering.

**Results:** CENTERED clustering revealed that ShA and LgA groups consisted of mixtures of different opioid user types.

**Conclusions:** CENTERED clustering revealed that experimenter-imposed grouping via differential access conditions limits our ability to identify distinct opioid user types that already exist naturally in the population.

## Introduction

Not all subjects with Opioid Use Disorders (OUD) benefit from pharmacotherapeutic treatment options (Bailey et al., 2025; Naji et al., 2025), suggesting that there are different types of opioid users. Indeed, current research indicates that individual patient characteristics, including genetic factors, comorbid conditions, and substance use history may play a significant role in the treatment response to FDA-approved buprenorphine or methadone (Hser et al., 2022, 2016; Kazi et al., 2024; Nielsen et al., 2022). Thus, to effectively address the OUD epidemic, it is crucial to understand these different opioid user types, as this knowledge can guide the development of more targeted and effective treatment strategies.

In preclinical research, opioid use paradigms have been developed to model different opioid user types. One model involves grouping rats self-administering drugs based on experimenter-determined drug exposure access time conditions into short-access (ShA) and long-access (LgA) groups. Evidence suggests that relative to ShA, LgA groups express behaviors akin to subjects that have developed substance use disorders, supporting face-validity for the idea that these groups represent different drug user types (Table 1). This approach assumes that ShA and LgA groups are behaviorally homogeneous within themselves and distinct from one another.

**Table 1:**
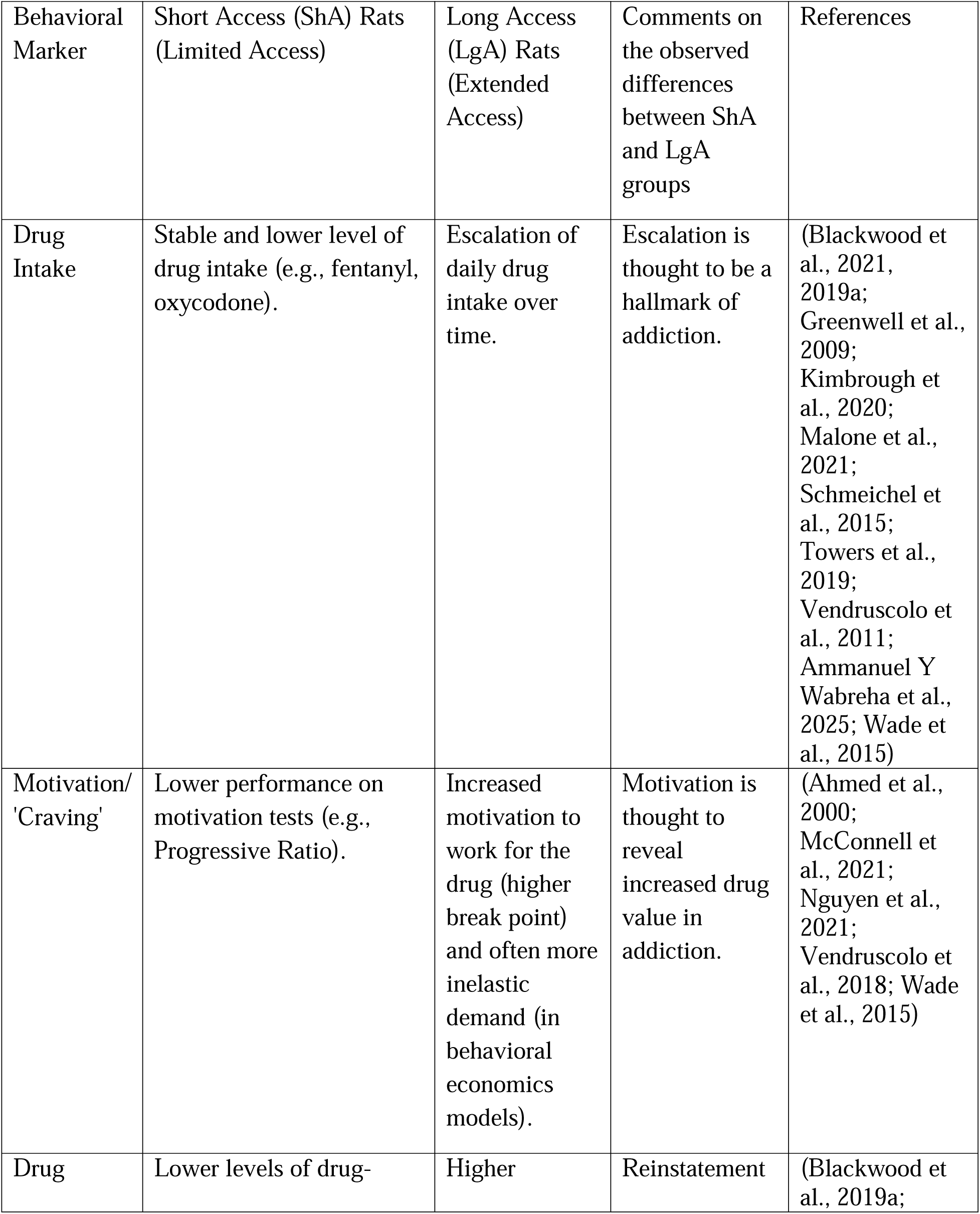
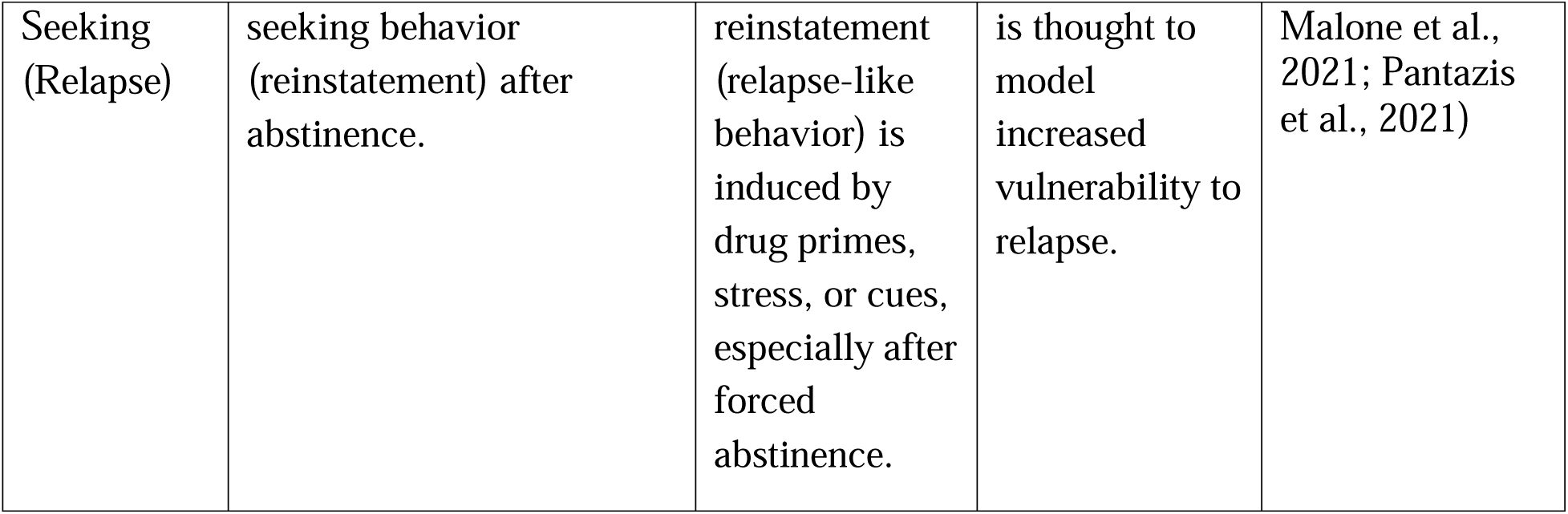
Short-access and Long-access groups may be different phenotypes of opioid users.

However, in all the above studies (Table 1), the experimenter decides, via random assignment, which subjects belong to which group (ShA versus LgA). Figure 1 proposes that for diverse populations there already exists heterogeneity *prior* to the experimenter-imposed grouping into ShA and LgA. Because the assignment of each subject into these groups is a random procedure, it is subjected to the laws of probability. In simple terms, if there were more than one type of opioid user in a population and we created groups by selecting individuals at random from this diverse population, the probability that the groups we create would be as diverse as the population is high. This means that if there were two types of individuals in the population, individuals from both types of opioid users would be represented in every group into which individuals are randomly assigned (Figure 1). Thus, ShA and LgA groups may not represent two distinct opioid user types, but instead mixtures of individuals belonging to different underlying phenotypes. If each group is composed of mixtures of distinct opioid user types, then comparing ShA and LgA as a way to understand distinct opioid user types is not ideal. It is important to confirm if this is the case as the whole access model rests on the idea that these groups are distinct.

**Figure 1:**
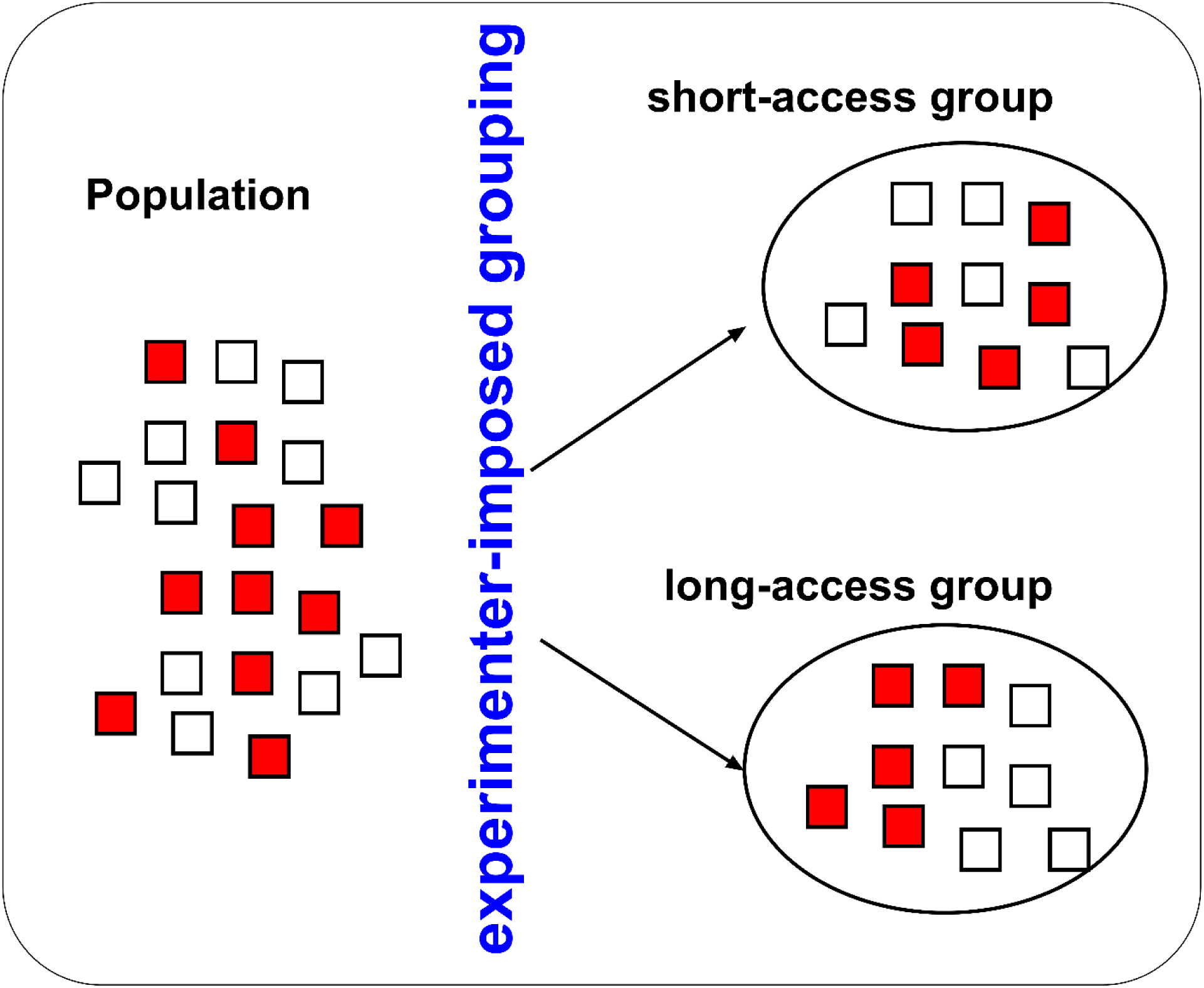
Proposed composition of groups containing randomly selected individuals (by experimenter) from a population composed of two drug user types. The above figure proposes that for diverse populations there is already heterogeneity *prior* to the experimenter-imposed grouping into short-access and long-access (ShA and LgA). The red and white squares represent individuals from two distinct drug user types *prior* to experimenter-imposed group assignment(s). Assuming the population contains a mix of individuals that belong to each opioid user type, the law of probability predicts that experimenter-assigned groups will consist of a mix of individuals from the different opioid user types. This implies that the assigned groups ShA and LgA should contain a mix of individuals from each type of opioid user. If this is so, the implication is that ShA versus LgA groups represent mixed groups of different opioid user types and are therefore not representative of distinct opioid user types, challenging the assumptions in the field.

The ShA will take less drug than the LgA not because they are necessarily less interested in the drug but because they have been restricted from taking more drug by the experimenter-imposed time constraint. The LgA will take more drug than the ShA not because they are necessarily more interested in the drug but because they are not as restricted as the ShA with regards to time for drug consumption. There may be individuals in the ShA group that are more motivated for the drug than some of the individuals in the LgA group, but the experimenter-assignment serves as an obstacle to identifying such individuals. To be able to understand the type(s) of opioid users in the population, we have to assess the drug use profile of each individual while controlling the differential access conditions. Thus, we require a new model that normalizes drug use as a function of drug experience and time. We developed the CENTERED (<u>C</u>umulative <u>E</u>xperience-<u>N</u>ormalized <u>T</u>ime-<u>E</u>ffect <u>R</u>esponse as an <u>E</u>xponential <u>D</u>ecay structure) model (Figure 2). CENTERED *clustering* is a procedure that involves principal component analysis-gaussian mixtures model *clustering* (PCA-GMM) of CENTERED model-derived variables to identify distinct groups of drug takers.

**Figure 2:**
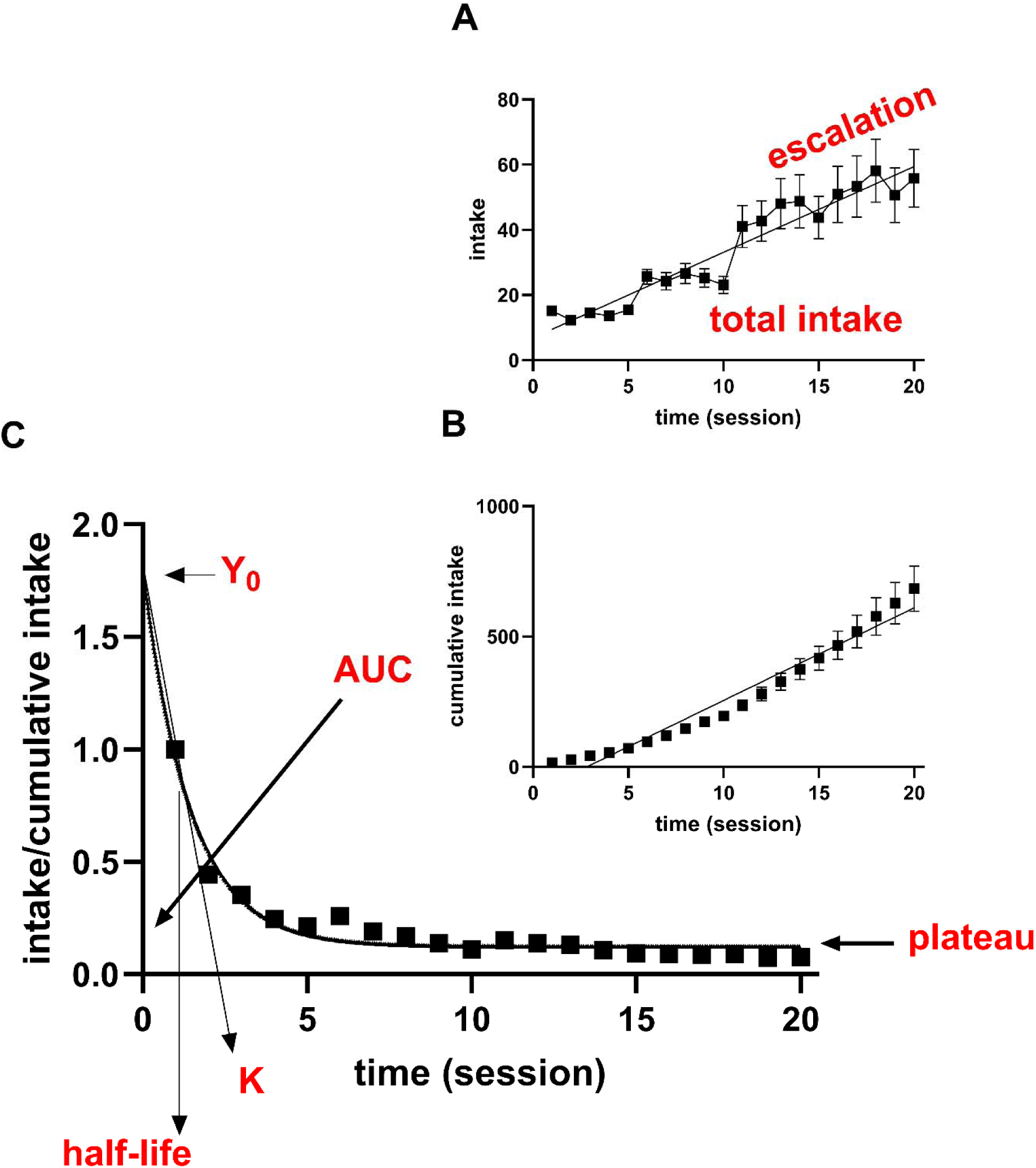
The CENTERED model. Graph A shows an intake-time graph (intake on the y-axis and time on the x-axis) to show the time course of drug intake (current model). From graph A we can obtain escalation (slope) using linear regression analysis and total intake as the sum of intake across time. Graph B shows a cumulative intake-time graph of the same data in graph A. The Cumulative Experience-Normalized Time-Effects on Response as an Exponential Decay structure (CENTERED) model is a plot of intake/cumulative intake on the y-axis versus time on the x-axis which is then fitted using an exponential decay curve function, see equation 1 to yield several variables including Y_0_ = intercept on the Y-axis at time = 0, minimum intake (plateau), exponential rate of change of intake (K), half-life (time to attain 50% of the Y_0_) and area under the curve (AUC) = Y_0_/K (equation 2). The CENTERED model accounts for drug intake as a function of cumulative intake and time.

Our hypothesis, based on Figure 1, is that groups assigned by the experimenter (ShA and LgA) consist of mixtures of individuals from distinct opioid user types identified via CENTERED clustering. To test our hypothesis, we reanalyzed behavioral data reported in (Blackwood et al., 2019a). Our methods, results and discussion are as below.

## Methods

The Methods are exactly as described in (Blackwood et al., 2019a). The rats in our study are the same as used in (Blackwood et al., 2019a). In that study, a total of thirty-eight (38) male Sprague Dawley rats (Charles River, Raleigh, NC, USA) were used. The rats were maintained on a 12-h reversed light/dark cycle and provided with food and water ad libitum. All procedures followed the guidelines outlined in the National Institutes of Health (NIH) Guide for the Care and Use of Laboratory Animals (eighth Edition) and were approved by the Animal Care and Use Committee for NIDA (National Institute of Drug Abuse). After rats were acclimated to their housing conditions, they were prepared for jugular catheter implantation surgery (Blackwood et al., 2019a).

Self-Administration experimental set up: Each chamber was equipped with two levers (active and inactive) located 8.5 cm above the grid floor. During experimentation, presses on the retractable active lever activated the infusion pump and tone-light cue whereas presses on the inactive lever had no reinforced consequences. Lever presses were reinforced using a fixed ratio-1 with a 20-s timeout accompanied by a 5-s compound tone-light cue. To connect the rats to the drug infusion system, we connected the catheter to the pump via polyethylene-50 tubing. To allow the rat to move freely, we connected this PE-50 tubing to a fluid swivel (Instech, Plymouth, PA). To preclude damage, the PE-50 tubing was protected by a metal spring. For details, see (Blackwood et al., 2019a).

Experimental Design: We allowed male Sprague Dawley rats to self-administer oxycodone (0.1 mg/kg/infusion, FR1, n = 30) or saline (n = 8) for 3h daily sessions for one week. Thereafter, we randomly grouped the rats into short access (ShA, 3h/session on week 2-4) and long access groups (LgA, 6h/session on week 2 and 9 h/session on week 3-4). For details, see (Blackwood et al., 2019a). We obtained intake data for d1-d20 (day 1 to day 20). For this study, we reanalyzed drug intake data obtained from the oxycodone takers (n = 30) and we did not analyze saline data (n = 8) because this study is focused on subjects with opioid experience.

### Current model grouping strategies

The current model is an example of experimenter-determined grouping strategy. The oxycodone subjects were grouped into ShA (n = 15) and LgA (n = 15) by the experimenter (Figure 1). The intake-time graphs of all individuals in the ShA and LgA groups are shown in Supplemental Figure 1 (Figure S1) and Figure S2, respectively.

### A new self-administration curve model – CENTERED model

The CENTERED model is an example of a Quantitative Structure Of Curve (QSOC) analytical model – it obtains variables that define the structure/shape of a curve. It is also an intake-experience-time curve: it is a combination of two curves (the intake-time curve – Figure 2A, and the cumulative-intake time curve – Figure 2B). CENTERED model curve is a plot of intake/cumulative intake on the y-axis versus time on the x-axis (Figure 2C). After plotting this curve, we obtained the variables that define the structure of the curve via the exponential decay fit equation (see below):

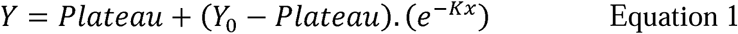

From this fit, we obtain the following variables for each individual: Y_0_ = intercept on the Y-axis at time = 0, minimum intake (plateau), exponential rate of change of intake (K), half-life (time to attain 50% of the Y_0_).

We also determined the area under the CENTERED curve (from zero to infinity) using the following function:

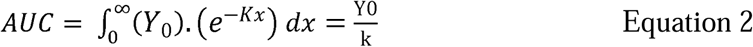

For each individual, we obtained intake-time curves (Figure 2A, individual graphs in Figure S1-2), cumulative intake-time curves (Figure 2B, individual graphs in Figure S3-4) and intake/cumulative intake-time curves (CENTERED model curve, Figure 2C, individual graphs in Figure S5-6).

From the intake-time curves (current model), we derived two (2) variables (escalation and total intake, Figure 2A). From the CENTERED model curve, we derived five (5) variables (Y_0_, plateau, K, half-life and AUC).

### A new drug user typology model: CENTERED clustering

After obtaining CENTERED model variables, we conducted <u>P</u>rincipal <u>C</u>omponent <u>A</u>nalysis-<u>G</u>aussian <u>M</u>ixtures <u>M</u>odel clustering (PCA-GMM) of these variables to determine if there were distinct clusters of individuals across all variables. We conducted dimensional reduction of the 5 variables using PCA to derive principal components and conducted GMM of these principal components to identify clusters. To ensure that we obtained the optimal number of clusters, we conducted 10 clustering/iterations and plotted a graph of Bayesian Information Criterion (BIC) versus the detected number of clusters (NCluster). The lowest BIC corresponds to the optimal number of clusters within the data set.

### Quantitative criteria for distinction of clusters

Clusters identified were determined to be distinct if they 1) have different centers with respect to principal component 1 (PC1, the principal component that can explain most of the variability in our data set), and 2) these groups should not be overlapping. Furthermore, we compared the clusters to determine if 1) they were distinct with regards to the averages of variables (using ANOVA), and 2) they were distinct with regards to the relationships between variables (using linear regression analysis).

### Model comparisons: Expected Results and Interpretation

We predicted that the CENTERED clustering of all model-related variables, irrespective of group assignment, would yield clusters that were distinct with regards to the above quantitative criteria and experimenter-assigned groups would include a mix of individuals from these distinct clusters.

Statistical Analysis: Statistical analysis was done using GraphPad Prism version 10 (GraphPad Software, San Diego, CA) and JMP Pro version 16 (SAS Institute Inc., Cary, NC). Alpha was set at 0.0001 for significance. We employed linear and non-linear (exponential decay) regression analysis to fit the time course curves for the intake-time and CENTERED model curve, respectively. We employed multivariate analysis to determine relationships between variables. We utilized PCA-GMM, regression analysis, and ANOVA to identify distinct clusters.

## Results

Graphs of individual intake-time curves, cumulative intake-time curves and CENTERED curves for oxycodone LgA and ShA groups are shown (appropriately labeled) in Supplemental Figure 1 (Figure S1) - Figure S6. For every individual, the variables (escalation and Total Intake) derived from the intake-time curves and the variables (Y_0_, plateau, K, half-life and AUC) derived from the CENTERED model curves for ShA and LgA are shown in Table S1-2, respectively.

### Self-administration curve model comparisons: CENTERED model versus current model (intake-time)

Multivariate analysis revealed that current model variables (escalation, Total Intake) are related to one of the CENTERED model variables (plateau) (Figure 3). Interestingly, the remaining CENTERED model variables are unrelated to the current model (Figure 3). Because most of the CENTERED model-derived variables were unrelated to current model-derived variables, we can confirm that these variables (and the model) are novel.

**Figure 3:**
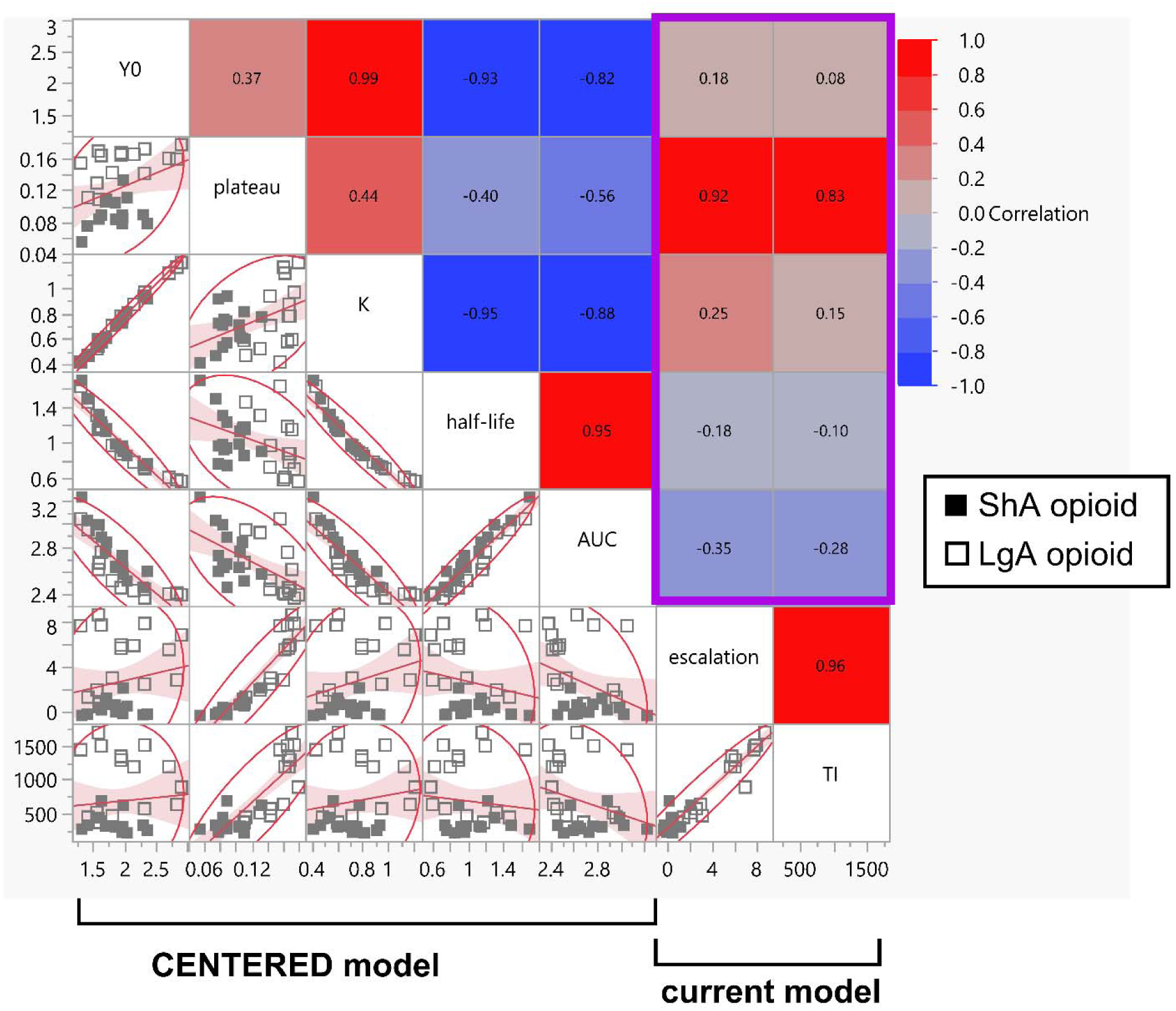
CENTERED model-derived variables are novel. For each individual (n = 30), current model variables (escalation, Total Intake) and CENTERED model variables (Y_0_, plateau, K, half-life, AUC) were obtained. We conducted multivariate analysis and report the correlation matrix above. We set significant difference at P = 0.0001. Escalation was linearly related to plateau (P < 0.0001) but unrelated to Y_0_ (P = 0.3374), K (P = 0.1818), half-life (P = 0.3371) and AUC (P = 0.0565). Total Intake was linearly related to plateau (P < 0.0001) but unrelated to Y_0_ (P = 0.6696), K (P = 0.4253), half-life (P = 0.5916) and AUC (P = 0.1388). The correlation coefficients are shown in the graph as heat maps. In summary, most of the variables revealed by the CENTERED model are not accounted for in the current model – these variables are novel.

### Drug User Typology Model comparisons: CENTERED clustering versus current model experimenter-determined grouping

We conducted PCA-GMM analysis on the CENTERED model-derived variables for all opioid users (n = 30) irrespective of what groups they were assigned to by the experimenter (ShA, LgA). PCA revealed a principal component 1 (PC 1) and PC 2 accounting for ∼80% and ∼16%, respectively, of all variability within the data set (Figure 4A). The variable loading on PC 1 (Figure 4B) revealed that Y_0_, K and plateau were positively related to PC 1 while half-life and AUC were negatively related to PC 1. Of all the variables, plateau was the least represented within PC 1 and the most represented in PC 2 (Figure 4B). The other variables contributed more to PC 1 than to PC 2. PCA-GMM identified 2 clusters. Figure 4C shows 2-D representation of identified clusters (including cluster composition as related to the experimenter-imposed groups) across PC 1 and PC 2. Figure 4D shows 3-D representation of identified clusters across PC 1-3. Figure 4E shows the optimal number of clusters for a plot of the BIC on the y-axis and the number of clusters identified on the x-axis. Figure 4F shows correlation matrix representation of identified clusters across the actual variables. Based on the quantitative criteria for distinction of clusters (see Methods section), these clusters were distinct across PC 1 and are non-overlapping (Figure 4C). Cluster1 included n = 18 subjects and cluster2 included n = 12 subjects.

**Figure 4:**
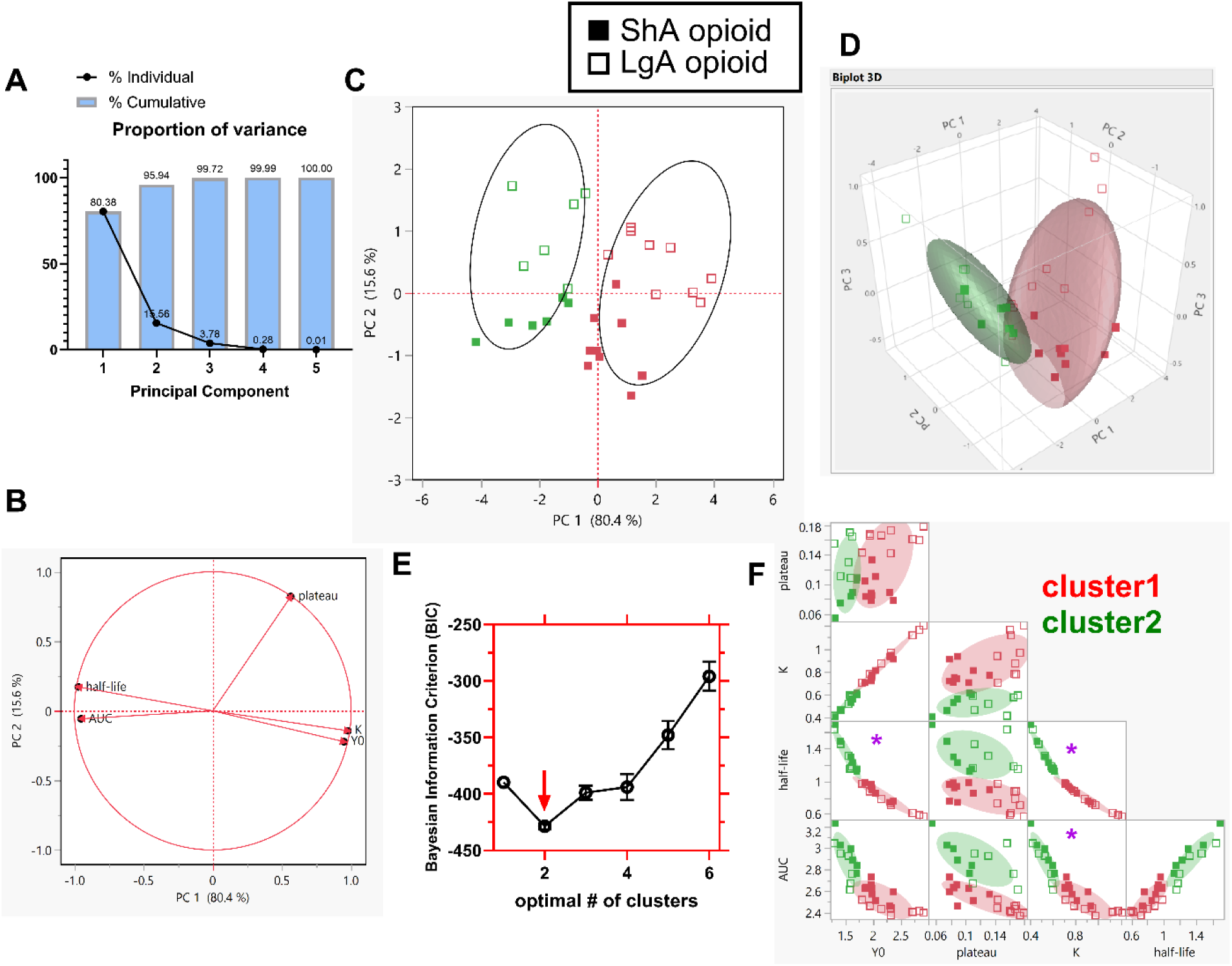
CENTERED clustering revealed two clusters of opioid user types: individuals from these two clusters were assigned into ShA and LgA groups by the experimenter. Principal component analysis- Gaussian mixtures model (PCA-GMM) clustering was conducted on CENTERED model variables for all oxycodone users (n = 30) irrespective of the groups defined by the experimenter. Graph A shows the proportion of variance represented by the principal components (PC) 1-4. Graph B shows the variable loading across PC 1 and PC 2. Graph C shows the clusters identified in a 2-D space defined by axes corresponding to PC 1- 2. Graph D shows the clusters identified in a 3-D space defined by axes corresponding to PC 1- 3. Graph E shows the optimum number of clusters from a plot of Bayesian Information Criterion (BIC) on the y-axis versus number of clusters on the x-axis – the optimum number of clusters = 2. Graph F shows the correlation matrix for actual variables showing the clusters identified. The closed boxes and open boxes represent individuals assigned to ShA and LgA groups, respectively. The red and green colors represent individuals in cluster1 and cluster2, respectively. CENTERED clustering revealed n = 18 rats in cluster1 and n = 12 rats in cluster2. These clusters were distinct with respect to PC 1 and did not overlap (Graph C). To the ShA group, the experimenter assigned n = 9 rats from cluster1 and n = 6 rats from cluster2. To the LgA group, the experimenter assigned n = 9 rats from cluster1 and n = 6 rats from cluster2. In summary, the experimenter-assigned groups consisted of a mixture of individuals from the two distinct clusters (opioid user types).

Interestingly, each experimenter-assigned group(s) included a mix of individuals from both of the clusters identified above. For example, the experimenter assigned n = 9 individuals from cluster1 and n = 6 individuals from cluster2 into the ShA group. Additionally, LgA group included n = 9 individuals from cluster1 and n = 6 individuals from cluster2. The composition of each cluster with regards to experimenter-imposed access time conditions are shown in Table 2.

**Table 2:** Group composition obtained via CENTERED clustering (A, cluster1 and cluster2) and current model clustering (B, cluster1 and cluster2) relative to groups assigned by experimenter (ShA, LgA).

|  | CENTERED clustering |  |
| --- | --- | --- |
| Experimenter-assigned group | Cluster1 | Cluster2 |
| ShA | N = 9 | N = 6 |
| LgA | N = 9 | N = 6 |
| total | N = 18 | N = 12 |

### Cluster1 versus Cluster2 were more behaviorally distinct than were ShA versus LgA

There were significant differences (P < 0.0001) between CENTERED model cluster1 and cluster2 for the slope of the relationship between variables Y_0_ and half-life, K and half-life, K and AUC (Table 3, column 2), but no significant differences in the slope of the relationship between other variables. For ShA versus LgA, we found no significant differences (P = 0.0001) with respect to the slopes of the relationship(s) between variables (Table 3, column 3).

**Table 3:** Comparisons of groups/clusters identified by three models (current model experimenter-determined grouping, current model clustering and CENTERED clustering) for significant differences (P = 0.0001) between the slope of the relationship between variables. Highlighted boxes show significant differences in slope of the relationships between the appropriately labeled relationships.

|  | Cluster1 v 2<br>(CENTERED<br>clustering) | ShA v LgA<br>(current model experimenter-determined grouping) | Batch1 v 2<br>(current model<br>clustering) |
| --- | --- | --- | --- |
| $Y_0$ and plateau | P = 0.8585 | P = 0.9514 | P = 0.3535 |
| $Y_0$ and K | P = 0.7668 | P = 0.2818 | P = 0.5963 |
| $Y_0$ and half-life | P < 0.0001 | P = 0.0122 | P = 0.6930 |
| $Y_0$ and AUC | P = 0.0074 | P = 0.0181 | P = 0.8702 |
| plateau and K | P = 0.2029 | P = 0.3645 | P = 0.7204 |
| plateau and half-life | P = 0.9335 | P = 0.8287 | P = 0.8568 |
| plateau and AUC | P = 0.6514 | P = 0.8767 | P = 0.8044 |
| K and half-life | P < 0.0001 | P = 0.0103 | P = 0.7408 |
| K and AUC | P < 0.0001 | P = 0.0096 | P = 0.8908 |
| half-life and AUC | P = 0.3209 | P = 0.1340 | P = 0.9140 |

We employed unpaired t-tests (with Welch’s correction) to determine if cluster1 and cluster2 were distinct with respect to all the CENTERED model-derived variables. There were differences between clusters for all (P < 0.0001, Figure 5A, C-E) except the plateau variable (P = 0.2824, Figure 5B). These groups were similar for 20% and different for 80% of the 5 variables compared.

**Figure 5:**
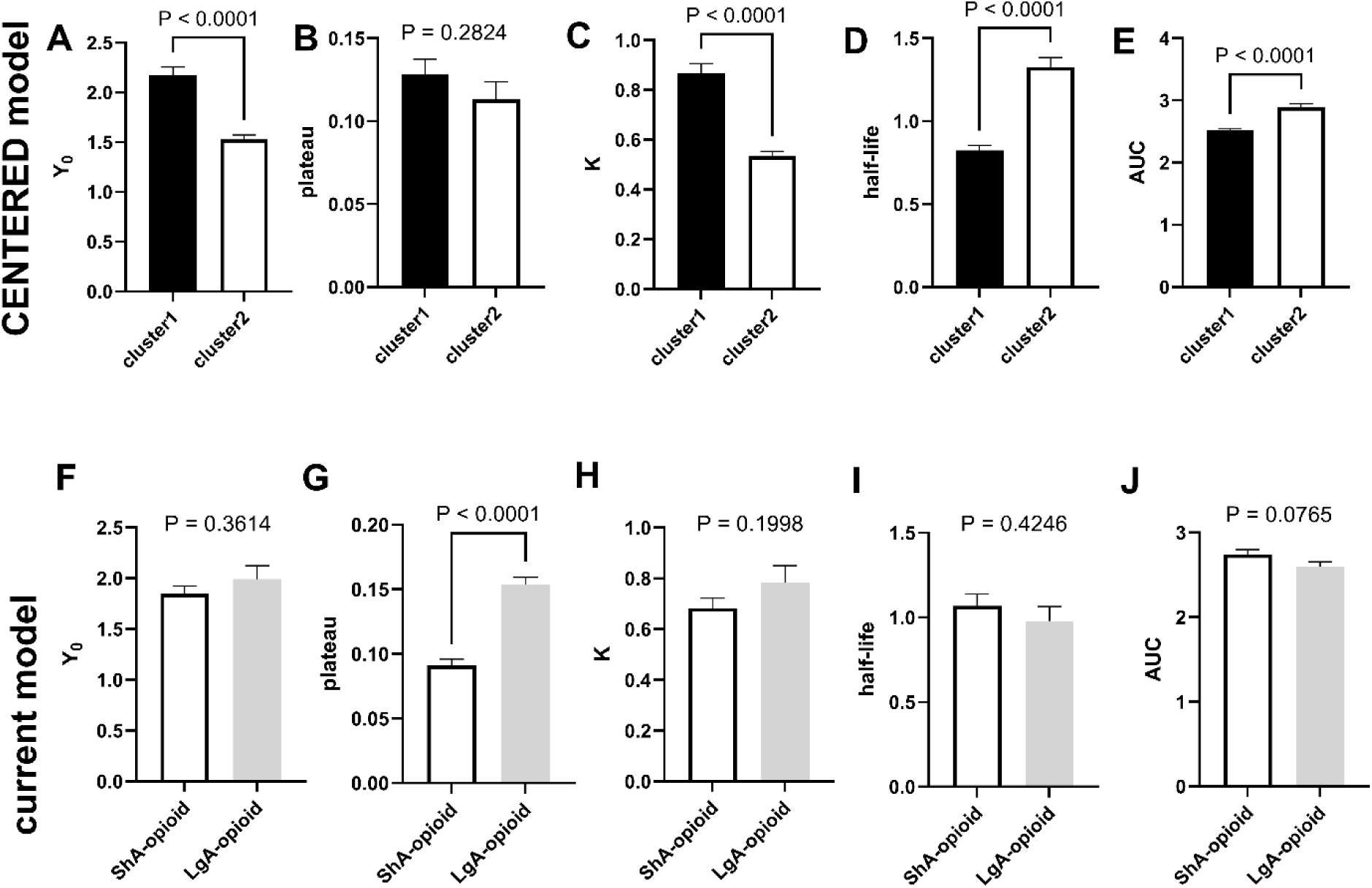
CENTERED clustering groups are more distinct than experimenter-assigned groups. CENTERED clustering revealed 2 groups. Unpaired t-tests (with Welch’s correction) revealed that these clusters were distinct (distinction set at P = 0.0001) with respect to all the CENTERED model variables, with the exception of plateau (Graphs A-E). In contrast, the experimenter-assigned groups of ShA and LgA were significantly different (P = 0.0001) ONLY for plateau and similar for all other CENTERED model variables (Graphs F-J). In summary, CENTERED clustering identified groups that were different for 80% of variables assessed and similar (not different) for 20% of variables assessed. Current model experimenter-defined groups were different for only 20% of the variables assessed and not different for 80% of the variables assessed. The CENTERED clustering identified clusters are more different from the groups assigned by the experimenter.

We employed unpaired t-tests (with Welch’s correction) to determine if ShA and LgA groups were distinct with respect to all the CENTERED model-derived variables. There were no differences between clusters for all variables (P > 0.0001, Figure 5F, H-J) except the plateau variable (P < 0.0001, Figure 5G).

### Drug self-administration intake-time course analysis: current model experimenter-determined grouping versus CENTERED clustering

At face value, LgA are distinct from ShA with respect to escalation and total intake (Figure 6A). Analysis of the intake-time curve using Two-way repeated measures ANOVA (with factors being experimenter-determined group – 2 levels and time – 20 levels) revealed the following: an experimenter-determined group × time interaction (F 2.958, 82.37 = 23.09, P < 0.0001), a main effect of experimenter-determined group (F 1, 28 = 31.25, P < 0.0001) and a main effect of time (F 2.958, 82.37 = 28.60, P < 0.0001). Linear regression analysis of the intake-time curve revealed differences between the escalation (F 1, 593 = 213, P < 0.0001). Unpaired t-tests (with Welch’s correction) showed significant differences between the two experimenter-determined groups with respect to average escalation (P < 0.0001, Figure 6B) and total intake (P < 0.0001, Figure 6C). With all these differences, the assumption would be that these belong to distinct drug user types based on the variables derived from the current model (intake-time curve).

**Figure 6:**
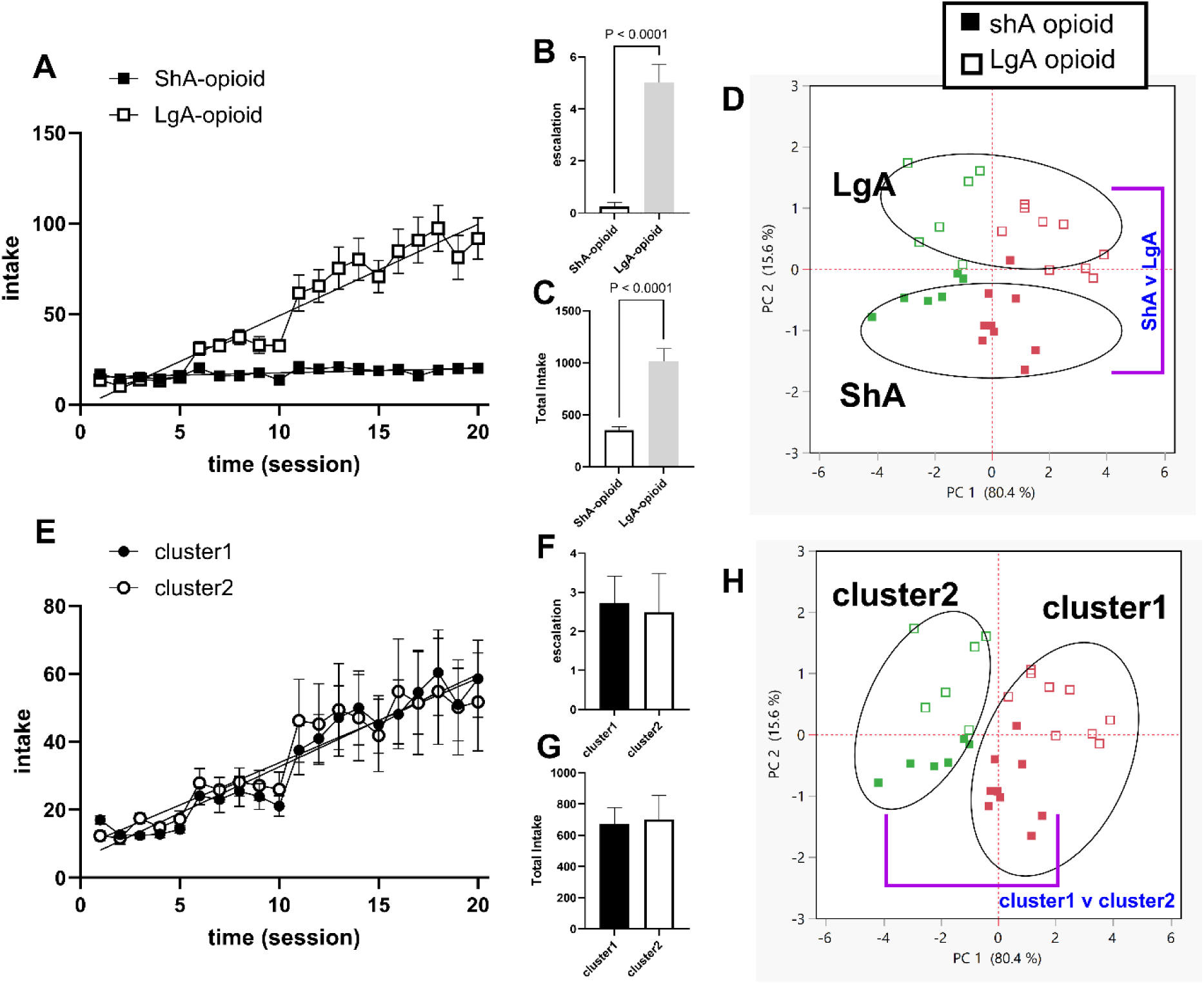
Escalation and total intake are not effective variables for distinguishing between different drug user types: importance of CENTERED clustering as a tool for drug user typology. Graph A shows the intake-time course of the groups defined by experimenter-assignment (ShA and LgA). Two-way repeated measures ANOVA and linear regression analysis revealed significant differences between these groups with respect to escalation and total intake (graphs A-C). However, when we accounted for CENTERED model variables and employed CENTERED clustering, these experimenter-defined groups were distinct with respect to PC 2 which corresponds to about 16% variability within the data set, but there were not distinct with regards to PC 1 which corresponds to about 80% variability within the data (graph D). On the other hand, the clusters identified via CENTERED clustering did not express differences in escalation and total intake (graphs E-G) yet were distinct with respect to 80% variability within the data set. Note that without the CENTERED model and CENTERED clustering this important information would remain unknown. In summary, groups expressing significant differences in escalation and total intake do not necessarily imply distinct groups. This has enormous implications for the field.

However, all these differences are only accounted for by about 16% of the variability within the data when we include variables from the new CENTERED model (see PC 2 axis in Figure 6D). We could clearly distinguish the experimenter-determined groups along the PC 2 axis (Figure 6D) but note that PC 2 ∼16% of the variability in the data set. This is in alignment with our determination that experimenter-determined groups were distinct only for 20% of the CENTERED model variables (Figure 5).

The clusters, on the other hand, are not distinct with respect to their intake-time curve (Figure 6E). For identified clusters, analysis of the intake-time curve using Two-way repeated measures ANOVA (with factors being cluster – 2 levels and time – 20 levels) revealed the following: no cluster × time interaction (F 1.677, 46.70 = 0.2635, P = 0.7306), no main effect of cluster (F 1, 28 = 0.01552, P = 0.9017), but a main effect of time (F 1.677, 46.70 = 14.81, P < 0.0001). Linear regression analysis of the intake-time curve revealed no differences between clusters for escalation variable (F 1, 593 = 0.2682, P = 0.6047). Unpaired t-tests (with Welch’s correction) showed no differences between cluster1 and cluster2 with respect to average escalation (P = 0.8485, Figure 6F) and total intake (P = 0.8752, Figure 6G). With the lack of any differences in the intake-time curve, one would assume that these clusters are the same.

However, the lack of differences are only relevant to about 20% of the variability within the data when we employ variables from the new CENTERED model. We could clearly distinguish clusters along the PC 1 axis (Figure 6H), and PC 1accounts for ∼80% of the variability in the data set. This is in alignment with our determination that experimenter-determined groups were distinct only for 20% of the CENTERED model variables (Figure 5).

### Current model clustering

We wanted to know if employing clustering analysis of escalation and total intake would yield distinct clusters (we term these as batches to avoid confusion with clusters identified via CENTERED clustering). We conducted PCA-GMM analysis on escalation and total intake data for all opioid users (n = 30) irrespective of what groups (ShA, LgA) they were assigned to by the experimenter. PCA revealed a principal component 1 (PC 1) and PC 2 accounting for ∼98% and ∼2%, respectively, of all variability within the data set (Figure 7A). The variable loading on PC 1 (Figure 7B) revealed that escalation and total intake were positively related to PC 1. Figure 7C shows 2-D representation of identified batches (including batch composition as related to the experimenter-imposed groups) across PC 1 and PC 2. Based on the quantitative criteria for distinction of clusters/batches (see Methods section), these batches are not clearly distinct across PC 1 though they are non-overlapping. Batch1 included n = 21 subjects and batch2 included n = 9 subjects. Interestingly, each experimenter-assigned group(s) (ShA and LgA) included a mix of individuals from both of the batches identified above. For example, to the ShA group, the experimenter randomly assigned n = 14 individuals from batch1 and n = 1 individual from batch2. To the LgA group, the experimenter assigned n = 7 individuals from batch1 and n = 8 individuals from batch2.

**Figure 7:**
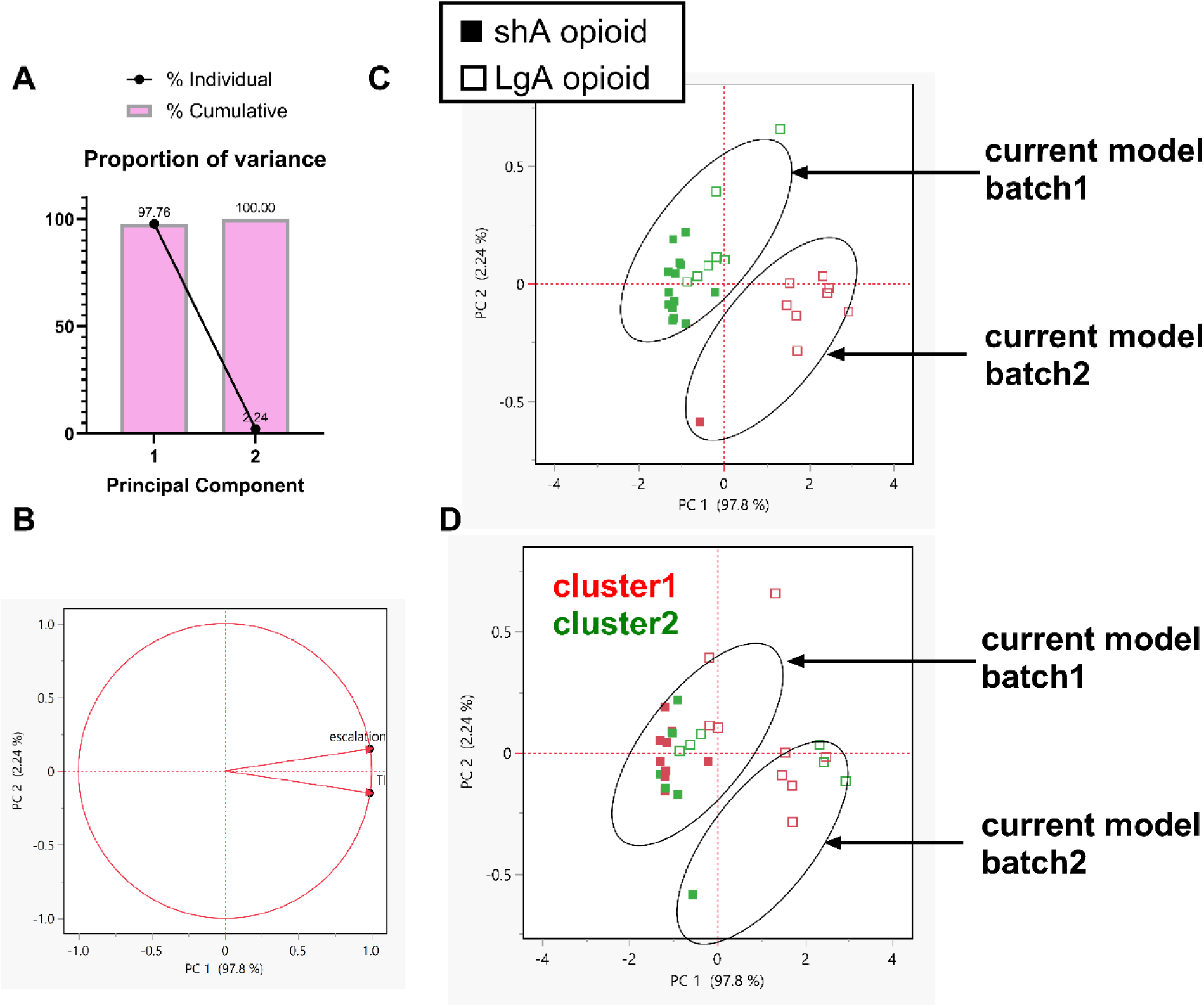
Clustering of escalation and total intake revealed batches different from the experimenter-assigned groups. The experimenter-assigned groups may be subjective groups. To check this, we conducted PCA-GMM clustering on escalation and total intake variables for all oxycodone users (n = 30) irrespective of the groups defined by the experimenter. Graph A shows the proportion of variance represented by the principal components (PC) 1-2. Graph B shows the variable loading across PC 1 and PC 2. Graph C shows the batches identified in a 2-D space defined by axes corresponding to PC 1- 2. Graph D is the same as C but shows the clusters (from CENTERED clustering) into which individuals belong. The closed and open boxes represent individuals assigned to ShA and LgA groups, respectively. The red and green colors represent cluster1 and cluster2, respectively. Current model clustering revealed n = 21 rats in batch1 and n = 9 rats in batch2. However, though the batches did not overlap, they were not distinct with respect to PC 1 (Graph C). Of the n = 21 rats identified as belonging to batch1, n = 14 rats were assigned (by the experimenter) to ShA group and n = 7 rats that were assigned to LgA groups. Of the n = 9 rats identified as belonging to batch2, n = 1 rat was assigned (by the experimenter) to ShA group and n = 8 rats were assigned to LgA groups. When we identified all individuals based on their identity via the CENTERED clustering, we found that of the n = 21 rats identified as belonging to batch1, n = 13 rats belonged to cluster1, and n = 8 rats belonged to cluster2. Furthermore, of the n = 9 rats identified as belonging to batch2, n = 5 rats belonged to cluster1, and n = 4 rats belonged to cluster2. Thus, clustering of escalation and total intake revealed batches of subjects that included a mix of individuals from distinct opioid user types. Importantly unbiased clustering did not reveal the same groups that the experimenter defined.

Note that the batches identified from clustering of escalation and total intake are not the same as the clusters identified from CENTERED clustering. Figure 7D is the same as Figure 7C but with members identified by the clusters they belonged to following CENTERED clustering. We showed earlier that there were n = 18 and n = 12 individuals in cluster1 and cluster2, respectively. Batch1 (n = 21) consists of a mix of individuals belonging to cluster1 (n = 13) and cluster2 (n = 8) whereas batch2 (n = 9) consists of a mix of individuals belonging to cluster1 (n = 5) and cluster2 (n = 4). While clustering escalation and total intake does not reveal clearly distinct clusters, when we identify individuals based on the (distinct) clusters they belong to following CENTERED clustering, the batches that emerge are mixtures of individuals from different clusters. With significant difference set at P = 0.0001, there were no differences between the slopes of the relationships between any of the CENTERED model-derived variables for the batches identified via current model clustering (Table 3, column 4).

## Discussion

It is generally assumed that subjects consuming opioids under differential access conditions represent distinct phenotypes (Table 1). However, based on the idea that heterogeneous populations already exist prior to experimenter-imposed assignments into groups based on differential access conditions (Figure 1), we hypothesized that experimenter-assigned groups would be composed of a mix of different opioid user types. In other words, individuals are *already* (potentially) behaviorally distinct before we assign them, albeit randomly, into ShA or LgA groups. These ShA and LgA groups will be composed of mixtures of individuals that represent different types of opioid users. To allow analysis of all subjects irrespective of differential access conditions we had to normalize our drug intake data to experience and time. Thus, we developed the CENTERED model (Figure 2). To test our hypothesis (using CENTERED clustering), we reanalyzed data from a previous study (Blackwood et al., 2019a) wherein we assigned oxycodone self-administering male rats into ShA and LgA. We confirmed that the variables revealed by the CENTERED model were novel (Figure 3). CENTERED clustering revealed two distinct clusters (two opioid user types) (Figure 4). The experimenter-assigned groups included a mix of individuals from the distinct clusters (Table 2). These clusters are more distinct from each other than ShA versus LgA groups (Figure 5-6, Table 3). Thus, we confirmed our hypothesis (Figure 1). This means that what we currently treat as a single experimental group (e.g., LgA) is not actually homogeneous but instead contains multiple underlying opioid user types. By confirming our hypothesis, we have revealed a major flaw in the current model that employs differential access conditions as a tool to understand different opioid user types - groups assigned randomly by the experimenter already consist of mixtures of distinct opioid user types.

While revealing the superiority of the CENTERED model over current models in identifying distinct drug users in the population, this study also reveals a weakness of the current intake-time curve model for drug user typology. Many current models distinguish drug user types based on how much drug they consume (total intake variable) or how they escalate their drug intake (escalation variable). Our findings suggest that groups identified using such variables may not represent truly distinct groups. Further, we determined that even when groups of subjects appear to be very distinct with respect to escalation and total intake (Figure 6A-C), they may be more similar than different when other (previously unknown) variables are also considered (Figure 6D). Conversely, groups of subjects may appear to be indistinguishable with respect to escalation and total intake variables (Figure 6E-G) but may be more different than similar (Figure 6H). While many studies have shown differences in escalation between ShA and LgA groups (Table 1, row 2), not all studies have observed this (Guha et al., 2022). Escalation may not be a sufficient variable for effective drug user typology.

Because of the limitations associated with experimenter-imposed grouping, we also conducted clustering of current model variables (escalation and total intake) of all subjects (Figure 7).

However, clustering did not clearly reveal batches that were similar to the experimenter-determined groups. Furthermore, these batches included a mix of individuals from both CENTERED model-identified clusters (opioid user types). The implication is that even with current models and current model-derived variables, the experimenter-assigned groups include a mix of individuals from very distinct clusters.

Unbiased clustering, which we employed in this study, addresses the limitations present in current approaches in the field (Table 4). In the clinical setting, unsupervised machine learning methods, including unbiased clustering analysis, have been employed to identify subtypes of opioid users (Afshar et al., 2019; Chu et al., 2023; Liu et al., 2023; Mullin et al., 2021; Mullin and Elkin, 2020; Panlilio et al., 2020; Shah-Mohammadi and Finkelstein, 2023). Unbiased clustering analysis has been successful in these studies in revealing hidden patterns of OUD behavior without the need for pre-defined categories, identifying patient factors affecting outcomes such as treatment success, abuse risk, and overdose risk. While these advancements in understanding opioid user types in the clinical settings have begun to be employed in preclinical settings (Allen et al., 2021; Kuhn et al., 2025a, 2025b), these new models are not yet widely utilized. To bridge the gaps between preclinical and clinical research, and to allow translational relevance of opioid user types identified in clinical and preclinical settings, there is a need to adopt unbiased data-driven models. Our work emphasizes this need.

**Table 4:**
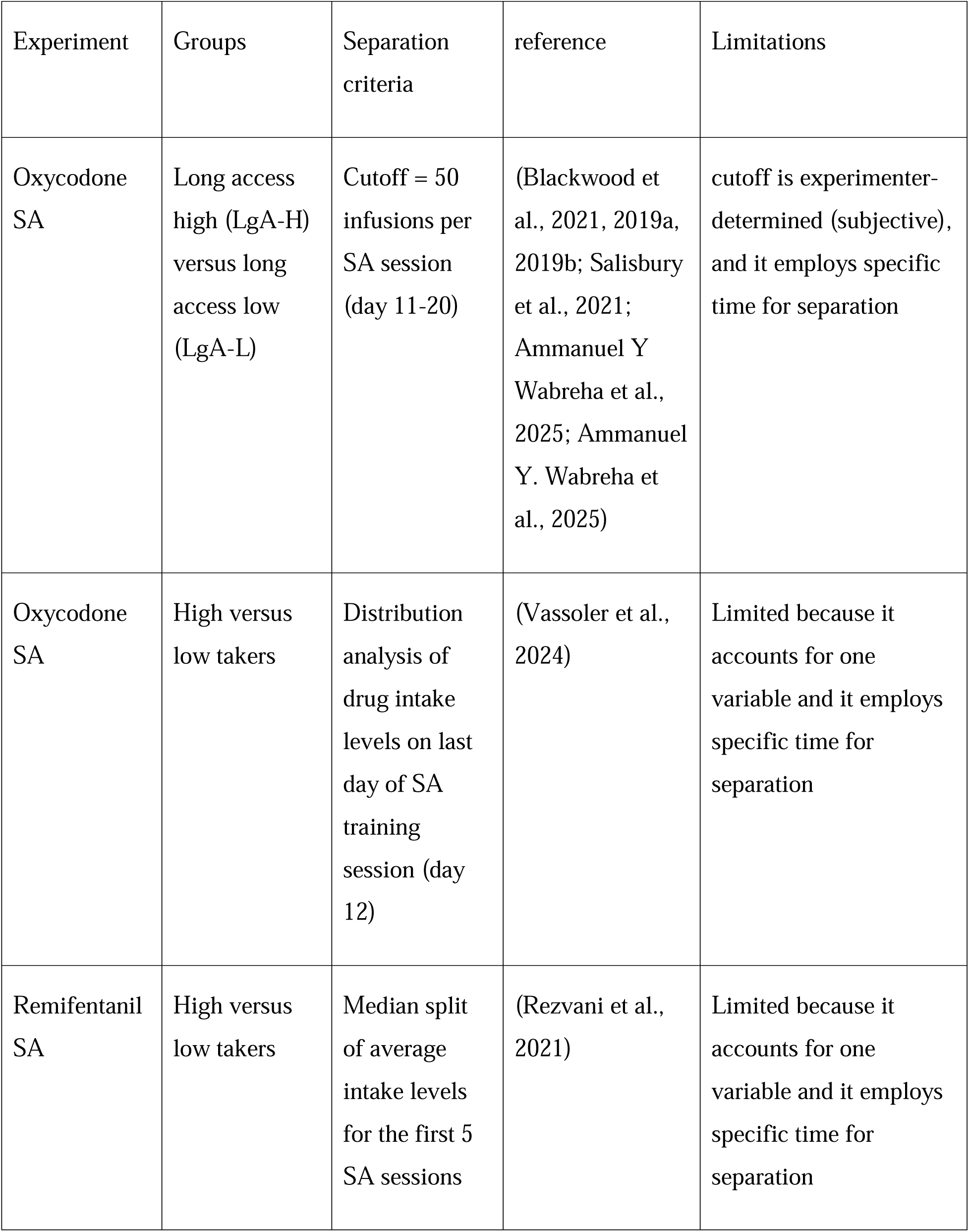
Limitations of current models that employ intake levels for identifying distinct opioid user types.

One limitation of this study is its generalizability, since female rats were not used. But this is because the study is entirely a re-analysis of already published data in which there were no female rats included (Blackwood et al., 2019a). That said, we have published evidence that differences between males and females in dopamine and psychostimulant-induced locomotor activity (Tigano and Job, 2025), and in methamphetamine self-administration are not due to biological sex. (Showell and Job, 2024), and as such we are confident that our findings with respect to males will be generalizable to females. That said, it will be important to verify this in future experiments that include female rats.

In summary, in this study, we present evidence challenging the assumption that experimenter-imposed grouping of subjects consuming opioids based on differential access conditions represent distinct phenotypes. We also show that variables obtained from the current intake-time curve model are not very effective for drug user typology. CENTERED clustering (a new approach) reveals the limitations of a current method and offers an alternative approach to move the field forward.

## Acknowledgement

The authors wish to acknowledge Dr Jean Lud Cadet, in whose lab the behavioral experiments were conducted. The authors wish to acknowledge Dr. Christopher Blackwood who was the first author of the study. The authors also wish to thank the other authors included in that study. The corresponding author of this manuscript was directly involved in all the behavioral experiments in the previous study.

---

**Figure S1:**
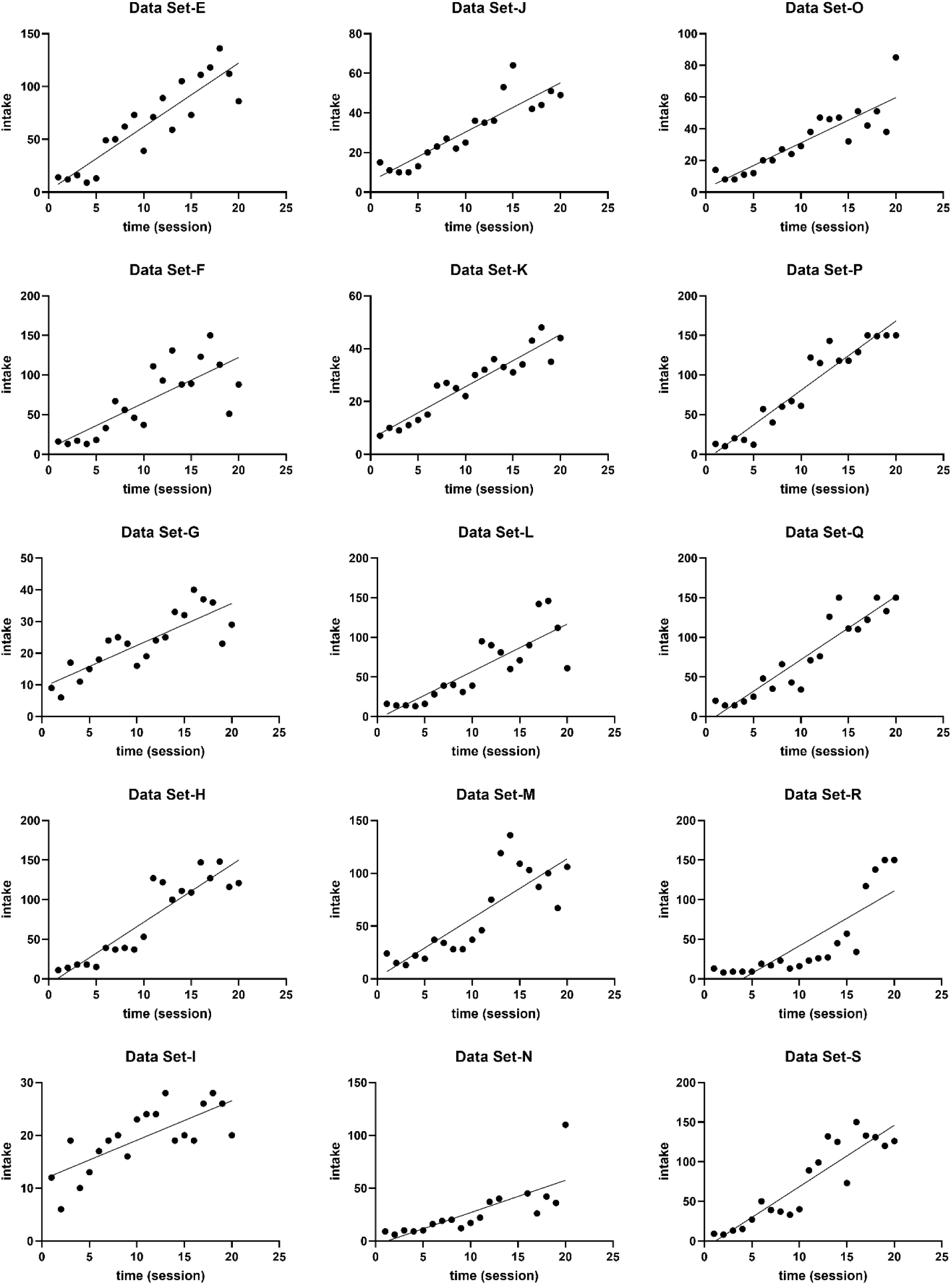
Individual intake-time graphs for all n = 15 oxycodone takers assigned to the long-access group.

**Figure S2:**
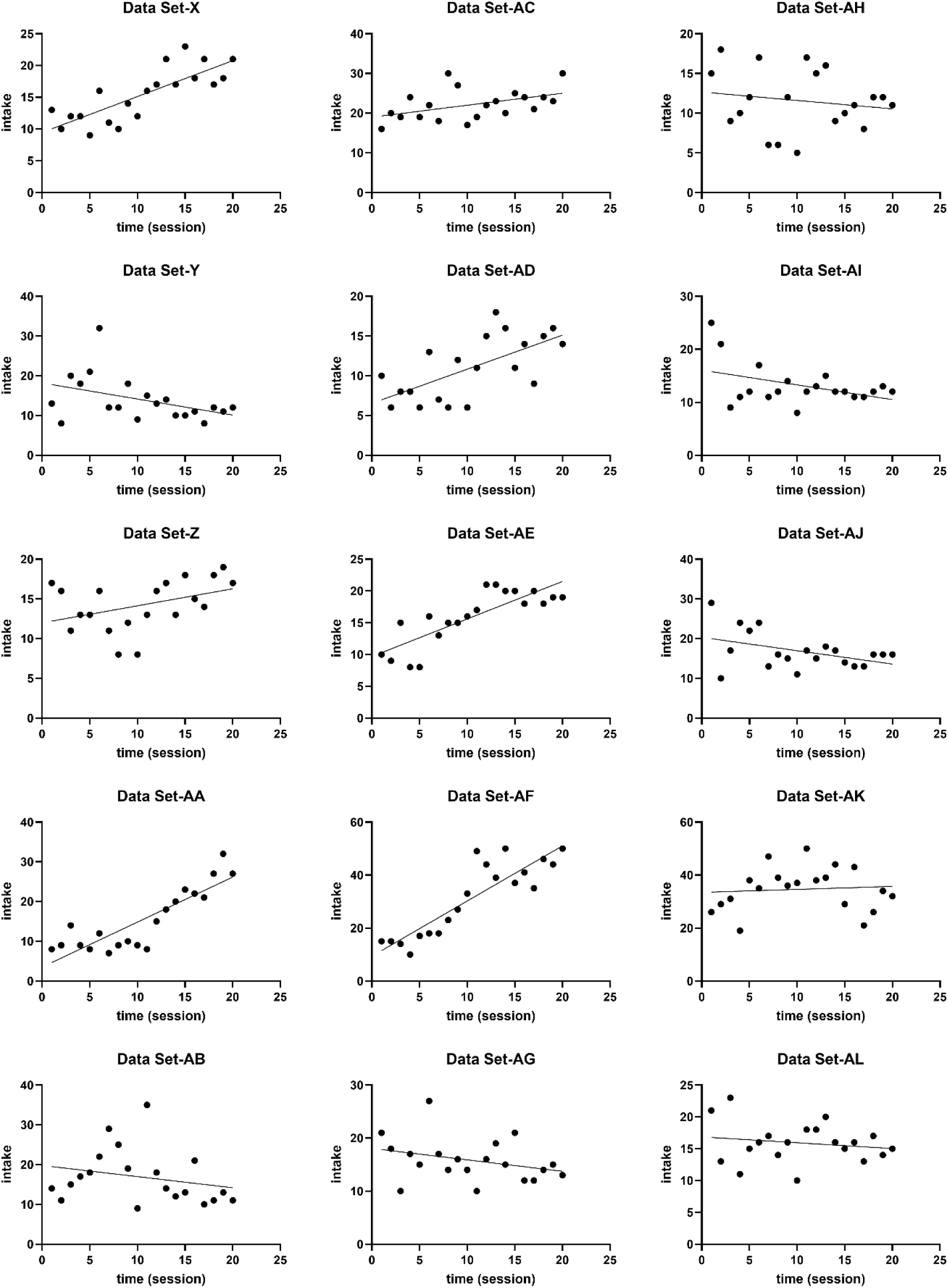
Individual intake-time graphs for all n = 15 oxycodone takers assigned to the short-access group.

**Figure S3:**
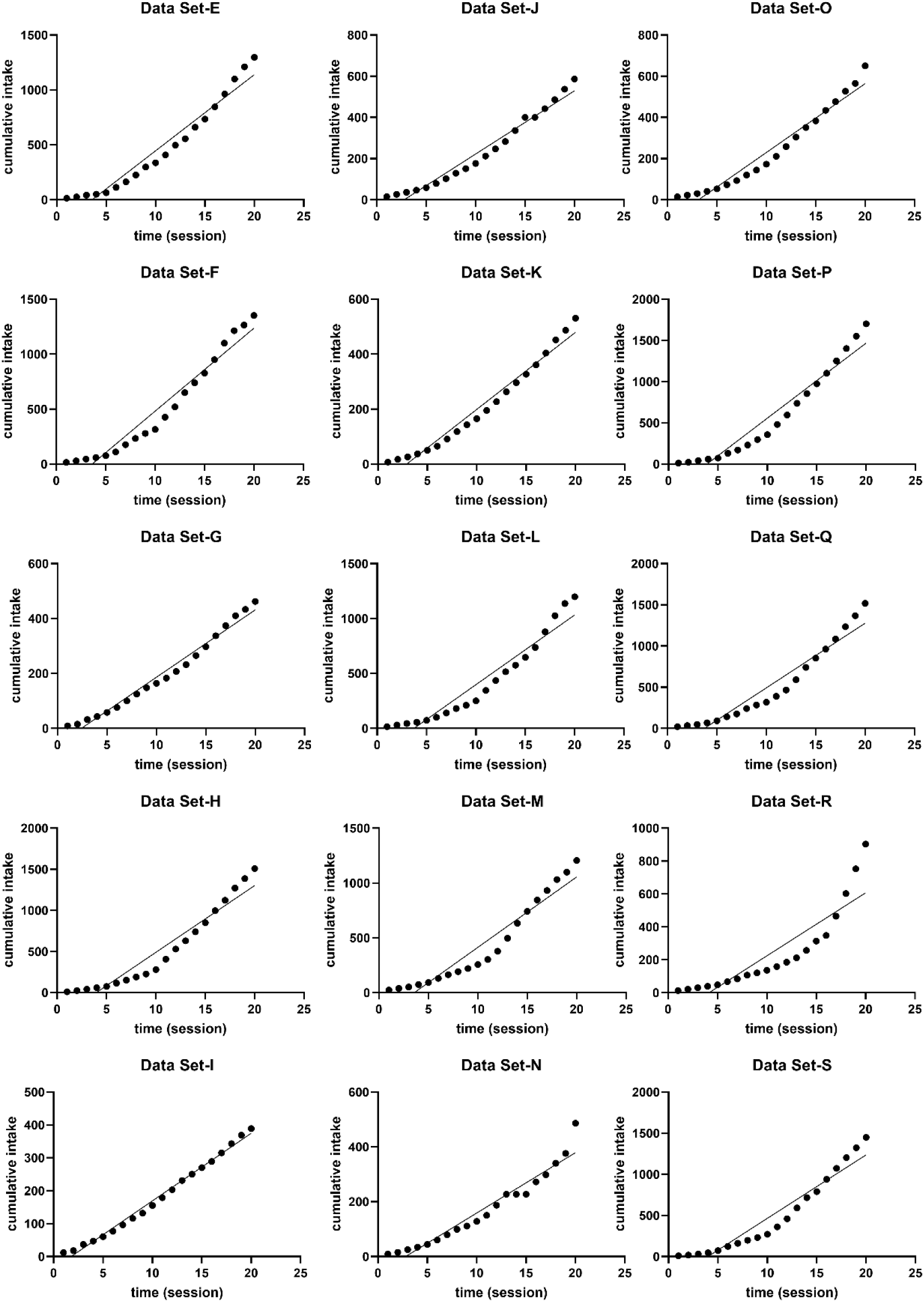
Individual cumulative intake-time data for all n = 15 oxycodone takers assigned to the long-access group.

**Figure S4:**
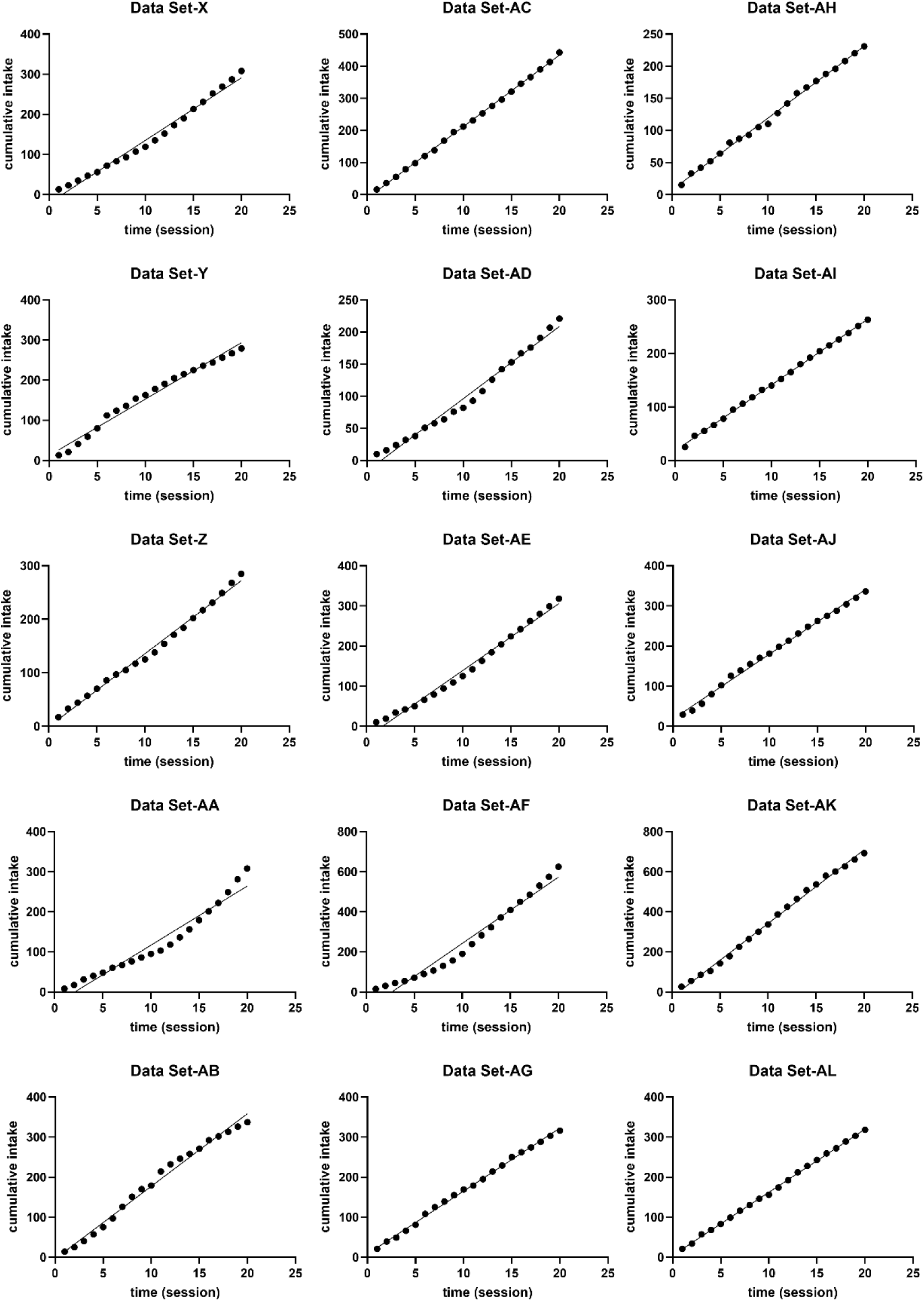
Individual cumulative intake-time data for all n = 15 oxycodone takers assigned to the short-access group.

**Figure S5:**
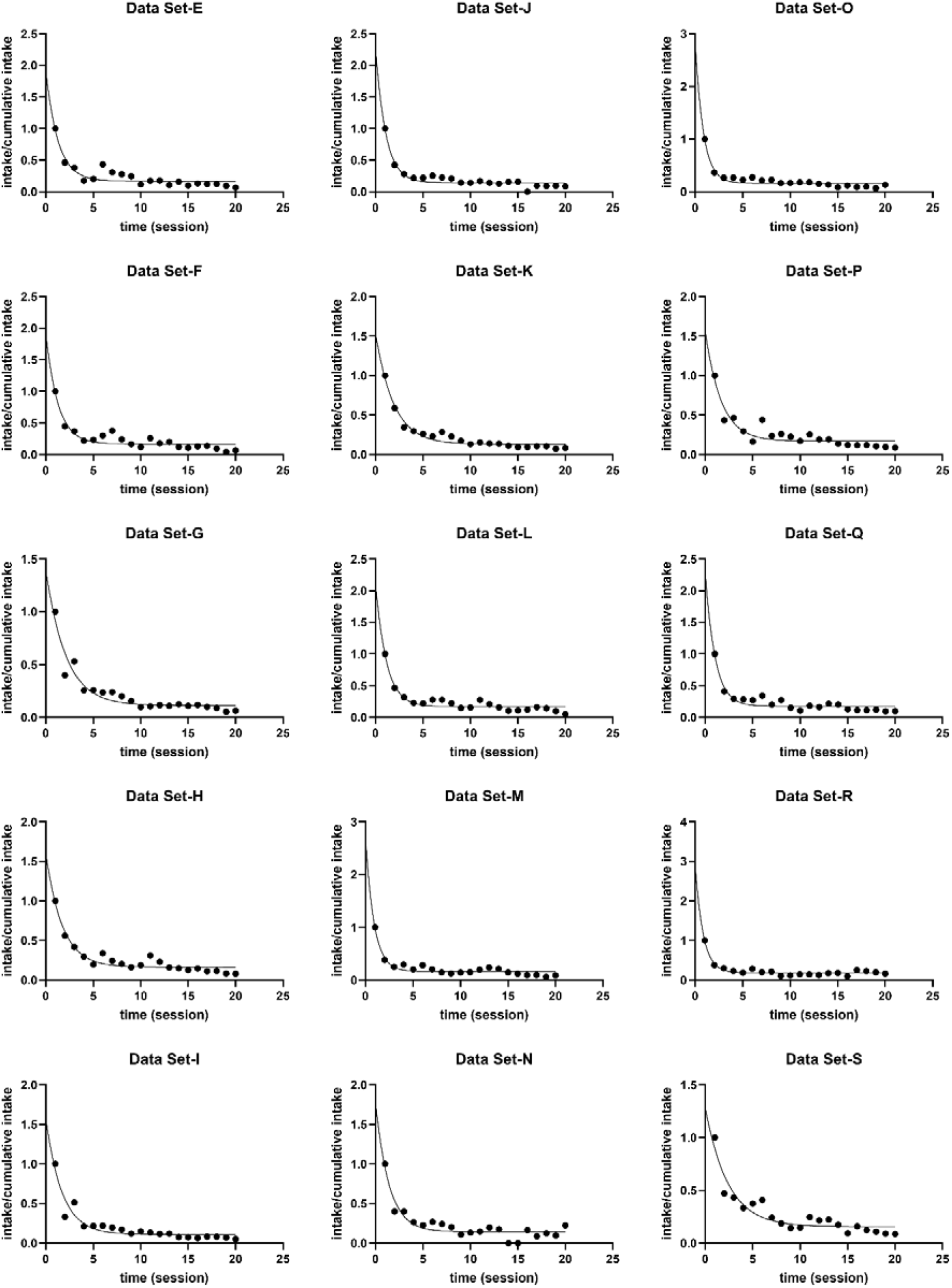
Individual intake/cumulative intake-time data for all n = 15 oxycodone takers assigned to the long-access group (CENTERED model).

**Figure S6:**
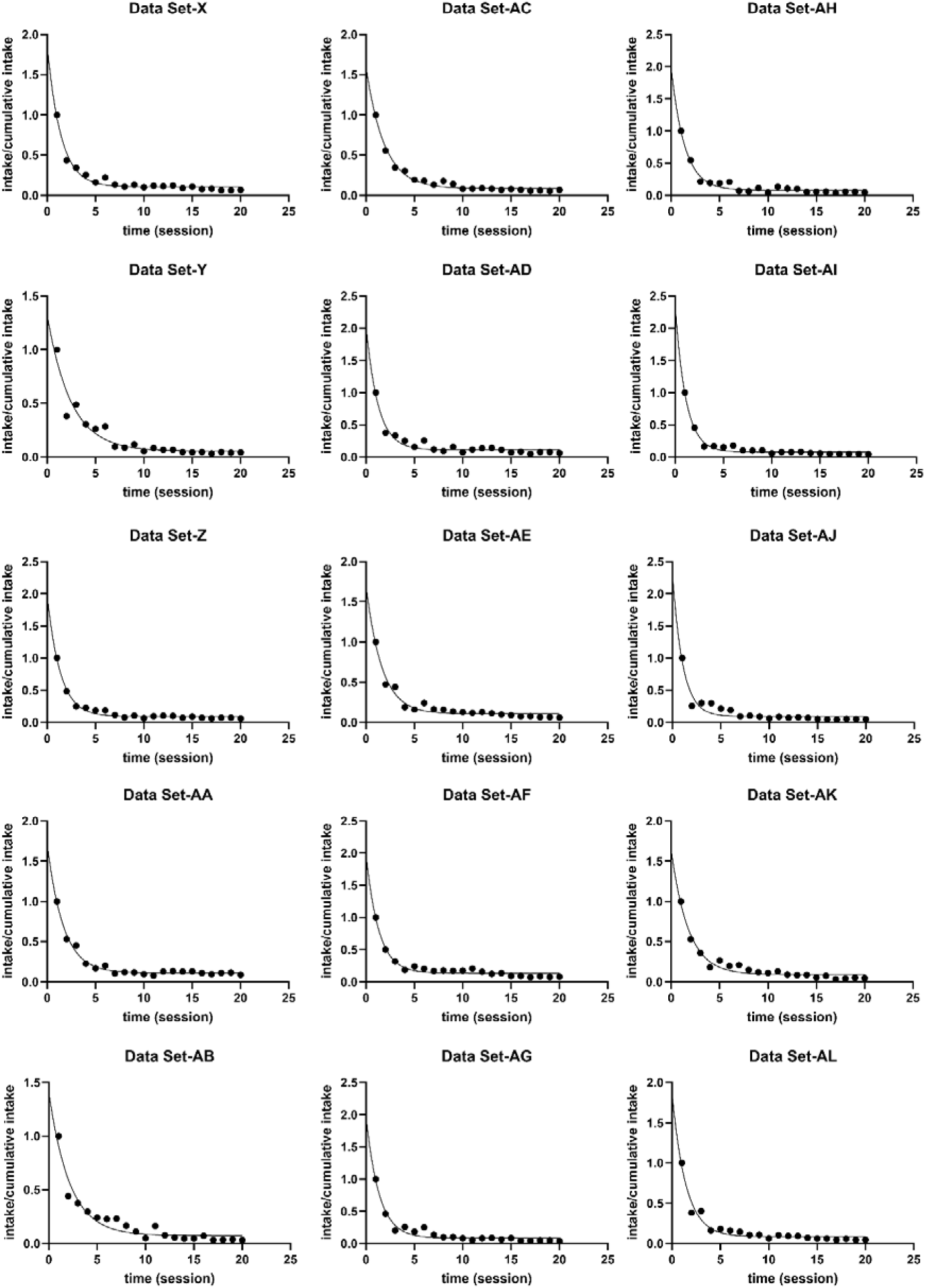
Individual intake/cumulative intake-time data for all n = 15 oxycodone takers assigned to the short-access group (CENTERED model).

**Table S1:** Current model (escalation, total intake TI) and CENTERED model variables (Y_0_, plateau, K, half-life and AUC) for all individuals grouped as long-access by the experimenter.

| ratID | escalation | Total intake | $Y_0$ | plateau | K | Half-life | AUC |
| --- | --- | --- | --- | --- | --- | --- | --- |
| E | 6.037 | 1297 | 1.935 | 0.1684 | 0.7838 | 0.8843 | 2.468742026 |
| F | 5.738 | 1353 | 1.934 | 0.1661 | 0.7851 | 0.8829 | 2.463380461 |
| G | 1.325 | 462 | 1.402 | 0.1116 | 0.4745 | 1.461 | 2.954689146 |
| H | 7.838 | 1509 | 1.613 | 0.165 | 0.5833 | 1.188 | 2.765300874 |
| I | 0.7466 | 389 | 1.586 | 0.1094 | 0.5928 | 1.169 | 2.675438596 |
| J | 2.487 | 586 | 2.307 | 0.1426 | 0.942 | 0.7358 | 2.449044586 |
| K | 1.975 | 531 | 1.546 | 0.1303 | 0.5273 | 1.315 | 2.931917315 |
| L | 5.988 | 1198 | 2.137 | 0.1668 | 0.876 | 0.7912 | 2.439497717 |
| M | 5.626 | 1205 | 2.702 | 0.1615 | 1.121 | 0.6182 | 2.410347904 |
| N | 3.061 | 486 | 1.792 | 0.1433 | 0.7097 | 0.9767 | 2.525010568 |
| O | 2.862 | 650 | 2.826 | 0.1601 | 1.168 | 0.5934 | 2.419520548 |
| P | 8.749 | 1702 | 1.572 | 0.1707 | 0.6004 | 1.154 | 2.618254497 |
| Q | 7.981 | 1517 | 2.309 | 0.1735 | 0.9719 | 0.7132 | 2.375758823 |
| R | 6.944 | 903 | 2.898 | 0.1788 | 1.206 | 0.5748 | 2.402985075 |
| S | 7.756 | 1449 | 1.289 | 0.1556 | 0.4228 | 1.639 | 3.0487228 |

**Table S2:** Current model (escalation, total intake TI) and CENTERED model variables (Y_0_, plateau, K, half-life and AUC) for all individuals grouped as short-access by the experimenter.

| ratID | escalation | Total intake | $Y_0$ | plateau | K | Half-life | AUC |
| --- | --- | --- | --- | --- | --- | --- | --- |
| X | 0.5654 | 308 | 1.855 | 0.1043 | 0.706 | 0.9817 | 2.627478754 |
| Y | -0.4053 | 279 | 1.319 | 0.05475 | 0.4085 | 1.697 | 3.228886169 |
| Z | 0.2143 | 285 | 1.967 | 0.08833 | 0.7402 | 0.9365 | 2.657389895 |
| AA | 1.132 | 308 | 1.711 | 0.1088 | 0.6024 | 1.151 | 2.840305445 |
| AB | -0.2805 | 337 | 1.408 | 0.07446 | 0.4648 | 1.491 | 3.029259897 |
| AC | 0.3015 | 443 | 1.595 | 0.08365 | 0.5331 | 1.3 | 2.991933971 |
| AD | 0.4293 | 221 | 2.042 | 0.1108 | 0.8125 | 0.8531 | 2.513230769 |
| AE | 0.591 | 318 | 1.702 | 0.1054 | 0.6155 | 1.126 | 2.765231519 |
| AF | 2.082 | 625 | 1.983 | 0.1328 | 0.7709 | 0.8992 | 2.57231807 |
| AG | -0.2165 | 316 | 1.975 | 0.0849 | 0.7511 | 0.9228 | 2.629476767 |
| AH | -0.106 | 231 | 1.968 | 0.07816 | 0.7207 | 0.9617 | 2.730678507 |
| AI | -0.2789 | 263 | 2.367 | 0.07804 | 0.9117 | 0.7603 | 2.596248766 |
| AJ | -0.3368 | 336 | 2.3 | 0.08906 | 0.9347 | 0.7416 | 2.460682572 |
| AK | 0.112 | 693 | 1.632 | 0.08886 | 0.5651 | 1.227 | 2.887984428 |
| AL | -0.09173 | 318 | 1.864 | 0.08355 | 0.7071 | 0.9802 | 2.636119361 |

## References

1. Afshar, M., Joyce, C., Dligach, D., Sharma, B., Kania, R., Xie, M., Swope, K., Salisbury-Afshar, E., Karnik, N.S., 2019. Subtypes in patients with opioid misuse: A prognostic enrichment strategy using electronic health record data in hospitalized patients. PLoS One 14. 10.1371/JOURNAL.PONE.0219717,

2. Ahmed, S.H., Walker, J.R., Koob, G.F., 2000. Persistent increase in the motivation to take heroin in rats with a history of drug escalation. Neuropsychopharmacology 22, 413–421. 10.1016/S0893-133X(99)00133-5

3. Allen, C., Kuhn, B.N., Cannella, N., Crow, A.D., Roberts, A.T., Lunerti, V., Ubaldi, M., Hardiman, G., Solberg Woods, L.C., Ciccocioppo, R., Kalivas, P.W., Chung, D., 2021. Network-Based Discovery of Opioid Use Vulnerability in Rats Using the Bayesian Stochastic Block Model. Front. Psychiatry 12.

4. Bailey, A.J., Votaw, V.R., Weiss, R.D., Mchugh, R.K., 2025. Capturing the Full Range of Buprenorphine Treatment Response. JAMA Psychiatry 82, 201–203. 10.1001/JAMAPSYCHIATRY.2024.3836,

5. Blackwood, C.A., Hoerle, R., Leary, M., Schroeder, J., Job, M.O., McCoy, M.T., Ladenheim, B., Jayanthi, S., Cadet, J.L., 2019a. Molecular Adaptations in the Rat Dorsal Striatum and Hippocampus Following Abstinence-Induced Incubation of Drug Seeking After Escalated Oxycodone Self-Administration. Mol. Neurobiol. 56, 3603–3615. 10.1007/S12035-018-1318-Z

6. Blackwood, C.A., Leary, M., Salisbury, A., McCoy, M.T., Cadet, J.L., 2019b. Escalated Oxycodone Self-Administration Causes Differential Striatal mRNA Expression of FGFs and IEGs Following Abstinence-Associated Incubation of Oxycodone Craving. Neuroscience 415, 173–183.

7. Blackwood, C.A., McCoy, M.T., Ladenheim, B., Cadet, J.L., 2021. Oxycodone self-administration activates the mitogen-activated protein kinase/ mitogen- and stress-activated protein kinase (MAPK-MSK) signaling pathway in the rat dorsal striatum. Sci. Rep. 11. 10.1038/S41598-021-82206-3

8. Chu, K., Carrière, G., Garner, R., Bosa, K., Hennessy, D., Sanmartin, C., 2023. Exploring the intersectionality of characteristics among those who experienced opioid overdoses: A cluster analysis. Heal. Reports 34, 3–14. 10.25318/82-003-X202300300001-ENG,

9. Greenwell, T.N., Walker, B.M., Cottone, P., Zorrilla, E.P., Koob, G.F., 2009. The α1 adrenergic receptor antagonist prazosin reduces heroin self-administration in rats with extended access to heroin administration. Pharmacol. Biochem. Behav. 91, 295–302. 10.1016/j.pbb.2008.07.012

10. Guha, S.K., Alonso-Caraballo, Y., Driscoll, G.S., Babb, J.A., Neal, M., Constantino, N.J., Lintz, T., Kinard, E., Chartoff, E.H., 2022. Ranking the contribution of behavioral measures comprising oxycodone self-administration to reinstatement of drug-seeking in male and female rats. Front. Behav. Neurosci. 16. 10.3389/FNBEH.2022.1035350/PDF

11. Hser, Y.I., Evans, E., Huang, D., Weiss, R., Saxon, A., Carroll, K.M., Woody, G., Liu, D., Wakim, P., Matthews, A.G., Hatch-Maillette, M., Jelstrom, E., Wiest, K., Mclaughlin, P., Ling, W., 2016. Long-term outcomes after randomization to buprenorphine/naloxone versus methadone in a multi-site trial. Addiction 111, 695–705. 10.1111/ADD.13238,

12. Hser, Y.I., Zhu, Y., Fei, Z., Mooney, L.J., Evans, E.A., Kelleghan, A., Matthews, A., Yoo, C., Saxon, A.J., 2022. Long-term follow-up assessment of opioid use outcomes among individuals with comorbid mental disorders and opioid use disorder treated with buprenorphine or methadone in a randomized clinical trial. Addiction 117, 151–161. 10.1111/ADD.15594,

13. Kazi, I., Chenoweth, M.J., Jutras-Aswad, D., Ahamad, K., Socias, M.E., Le Foll, B., Tyndale, R.F., 2024. Pharmacogenetics of Biochemically Verified Abstinence in an Opioid Agonist Therapy Randomized Clinical Trial of Methadone and Buprenorphine/Naloxone. Clin. Pharmacol. Ther. 115, 506–514. 10.1002/CPT.3112,

14. Kimbrough, A., Kononoff, J., Simpson, S., Kallupi, M., Sedighim, S., Palomino, K., Conlisk, D., Momper, J.D., de Guglielmo, G., George, O., 2020. Oxycodone self-administration and withdrawal behaviors in male and female Wistar rats. Psychopharmacology (Berl). 237, 1545–1555. 10.1007/S00213-020-05479-Y,

15. Kuhn, B.N., Cannella, N., Chitre, A.S., Nguyen, K.M.H., Cohen, K., Chen, D., Peng, B., Ziegler, K.S., Lin, B., Johnson, B.B., Missfeldt Sanches, T., Crow, A.D., Lunerti, V., Gupta, A., Dereschewitz, E., Soverchia, L., Hopkins, J.L., Roberts, A.T., Ubaldi, M., Abdulmalek, S., Kinen, A., Hardiman, G., Chung, D., Polesskaya, O., Solberg Woods, L.C., Ciccocioppo, R., Kalivas, P.W., Palmer, A.A., 2025a. Genome-wide association study reveals multiple loci for nociception and opioid consumption behaviors associated with heroin vulnerability in outbred rats. Mol. Psychiatry 30, 3363–3375. 10.1038/S41380-025-02922-4,

16. Kuhn, B.N., Cannella, N., Crow, A.D., Lunerti, V., Gupta, A., Walterhouse, S.J., Allen, C., Chalhoub, R.M., Dereschewitz, E., Roberts, A.T., Cockerham, M., Beeson, A., Nall, R.W., Palmer, A.A., Hardiman, G., Solberg Woods, L.C., Chung, D., Ciccocioppo, R., Kalivas, P.W., 2025b. Distinct Behavioral Profiles and Neuronal Correlates of Heroin Vulnerability Versus Resiliency in a Multi-Symptomatic Model of Heroin Use Disorder in Rats. Am. J. Psychiatry 182, 198–208. 10.1176/APPI.AJP.20230623,

17. Liu, P., Wang, Z., Liu, N., Peres, M.A., 2023. A scoping review of the clinical application of machine learning in data-driven population segmentation analysis. J. Am. Med. Informatics Assoc. 30, 1573–1582. 10.1093/JAMIA/OCAD111,

18. Malone, S.G., Keller, P.S., Hammerslag, L.R., Bardo, M.T., 2021. Escalation and reinstatement of fentanyl self-administration in male and female rats. Psychopharmacology (Berl). 238, 2261–2273. 10.1007/S00213-021-05850-7,

19. McConnell, S.A., Brandner, A.J., Blank, B.A., Kearns, D.N., Koob, G.F., Vendruscolo, L.F., Tunstall, B.J., 2021. Demand for fentanyl becomes inelastic following extended access to fentanyl vapor self-administration. Neuropharmacology 182. 10.1016/j.neuropharm.2020.108355

20. Mullin, S., Elkin, P., 2020. Assessing opioid use patient representations and subtypes. Stud. Health Technol. Inform. 270, 823–827. 10.3233/SHTI200276,

21. Mullin, S., Zola, J., Lee, R., Hu, J., MacKenzie, B., Brickman, A., Anaya, G., Sinha, S., Li, A., Elkin, P.L., 2021. Longitudinal K-means approaches to clustering and analyzing EHR opioid use trajectories for clinical subtypes. J. Biomed. Inform. 122. 10.1016/j.jbi.2021.103889

22. Naji, L., Rosic, T., Dennis, B., Worster, A., Paul, J., Thabane, L., Samaan, Z., 2025. Effectiveness of methadone versus buprenorphine in the treatment of opioid use disorder: secondary analyses of prospective cohort study data. BMJ Open 15. 10.1136/BMJOPEN-2024-095645,

23. Nguyen, J.D., Grant, Y., Taffe, M.A., 2021. Paradoxical changes in brain reward status during oxycodone self-administration in a novel test of the negative reinforcement hypothesis. Br. J. Pharmacol. 178, 3797–3812. 10.1111/BPH.15520

24. Nielsen, S., Tse, W.C., Larance, B., 2022. Opioid agonist treatment for people who are dependent on pharmaceutical opioids. Cochrane Database Syst. Rev. 2022. 10.1002/14651858.CD011117.PUB3,

25. Panlilio, L. V., Stull, S.W., Bertz, J.W., Burgess-Hull, A.J., Kowalczyk, W.J., Phillips, K.A., Epstein, D.H., Preston, K.L., 2020. Beyond abstinence and relapse: cluster analysis of drug-use patterns during treatment as an outcome measure for clinical trials. Psychopharmacology (Berl). 237, 3369–3381. 10.1007/S00213-020-05618-5,

26. Pantazis, C.B., Gonzalez, L.A., Tunstall, B.J., Carmack, S.A., Koob, G.F., Vendruscolo, L.F., 2021. Cues conditioned to withdrawal and negative reinforcement: Neglected but key motivational elements driving opioid addiction. Sci. Adv. 7. 10.1126/SCIADV.ABF0364

27. Rezvani, A.H., Wells, C., Hawkey, A., Blair, G., Koburov, R., Ko, A., Schwartz, A., Kim, V.J., Levin, E.D., 2021. Differential behavioral functioning in the offspring of rats with high vs. low self-administration of the opioid agonist remifentanil. Eur. J. Pharmacol. 909. 10.1016/j.ejphar.2021.174407

28. Salisbury, A.J., Blackwood, C.A., Cadet, J.L., 2021. Prolonged Withdrawal From Escalated Oxycodone Is Associated With Increased Expression of Glutamate Receptors in the Rat Hippocampus. Front. Neurosci. 14. 10.3389/FNINS.2020.617973,

29. Schmeichel, B.E., Barbier, E., Misra, K.K., Contet, C., Schlosburg, J.E., Grigoriadis, D., Williams, J.P., Karlsson, C., Pitcairn, C., Heilig, M., Koob, G.F., Vendruscolo, L.F., 2015. Hypocretin receptor 2 antagonism dose-dependently reduces escalated heroin self-administration in rats. Neuropsychopharmacology 40, 1123–1129. 10.1038/NPP.2014.293

30. Shah-Mohammadi, F., Finkelstein, J., 2023. Identification of Subphenotypes of Opioid Use Disorder Using Unsupervised Machine Learning. Stud. Health Technol. Inform. 302, 897–898. 10.3233/SHTI230299,

31. Showell, B.H., Job, M.O., 2024. The most significant differences between male and female rats regarding psychostimulant self-administration behavior are unrelated to biological sex.

32. Tigano, A.M., Job, M.O., 2025. Lack of Sex Differences in Psychostimulant-Induced Locomotor Activity When Comparing Rats From the Same Behavioral Groups. Biol. Psychiatry Glob. Open Sci. 5. 10.1016/j.bpsgos.2025.100519

33. Towers, E.B., Tunstall, B.J., McCracken, M.L., Vendruscolo, L.F., Koob, G.F., 2019. Male and female mice develop escalation of heroin intake and dependence following extended access. Neuropharmacology 151, 189–194. 10.1016/J.NEUROPHARM.2019.03.019

34. Vassoler, F.M., Budge, K.E., Isgate, S.B., Gildawie, K.R., Byrnes, E.M., 2024. Neuroplasticity-related genes correlate with individual differences in distinct phases of oxycodone self-administration in male rats. Neuropharmacology 254. 10.1016/j.neuropharm.2024.109972

35. Vendruscolo, J.C.M., Tunstall, B.J., Carmack, S.A., Schmeichel, B.E., Lowery-Gionta, E.G., Cole, M., George, O., Vandewater, S.A., Taffe, M.A., Koob, G.F., Vendruscolo, L.F., 2018. Compulsive-Like Sufentanil Vapor Self-Administration in Rats. Neuropsychopharmacology 43, 801–809. 10.1038/NPP.2017.172

36. Vendruscolo, L.F., Schlosburg, J.E., Misra, K.K., Chen, S.A., Greenwell, T.N., Koob, G.F., 2011. Escalation patterns of varying periods of heroin access. Pharmacol. Biochem. Behav. 98, 570–574. 10.1016/J.PBB.2011.03.004

37. Wabreha, Ammanuel Y, Adjei, N., Ladenheim, B., Cadet, J.L., Daiwile, A.P., 2025. Escalated oxycodone self-administration is associated with expression of voltage gated and calcium activated potassium channels in the mesocorticolimbic system in rats. Front. Pharmacol. 16, 1653356. 10.3389/fphar.2025.1653356

38. Wabreha, Ammanuel Y., McCoy, M.T., Cadet, J.L., Daiwile, A.P., 2025. Escalated Oxycodone Self-Administration Is Associated with Activation of Specific Gene Networks in the Rat Dorsal Striatum. Int. J. Mol. Sci. 26, 7356.

39. Wade, C.L., Vendruscolo, L.F., Schlosburg, J.E., Hernandez, D.O., Koob, G.F., 2015. Compulsive-like responding for opioid analgesics in rats with extended access. Neuropsychopharmacology 40, 421–428. 10.1038/NPP.2014.188

